# Lysosomal degradation ensures accurate chromosomal segregation to prevent genomic instability

**DOI:** 10.1101/802025

**Authors:** Eugènia Almacellas, Charles Day, Santiago Ambrosio, Albert Tauler, Caroline Mauvezin

## Abstract

Lysosomes, as primary degradative organelles, are the end-point of different converging pathways including macroautophagy. To date, lysosome function has mainly focused on interphase cells, while their role during mitosis remains controversial. Mitosis dictates the faithful transmission of genetic material among generations, and perturbations of mitotic division lead to chromosomal instability, a hallmark of cancer. Heretofore, correct mitotic progression relies on the orchestrated degradation of mitotic factors, which was mainly attributed to ubiquitin-triggered proteasome-dependent degradation. Here, we show that mitotic transition does not only rely on proteasome-dependent degradation, as impairment of lysosomes increases mitotic timing and leads to mitotic errors, thus promoting chromosomal instability. Furthermore, we identified several putative lysosomal targets in mitotic cells. Among them, WAPL, a cohesin regulatory protein, emerged as a novel p62-interacting protein for targeted lysosomal degradation. Finally, we characterized an atypical nuclear phenotype, the toroidal nucleus, as a novel biomarker for genotoxic screenings. Our results establish lysosome-dependent degradation as an essential event to prevent genomic instability.

## INTRODUCTION

Chromosomal instability (CIN) is defined as an abnormal loss or rearrangement of chromosomes during cell division^1^ and positively correlates with poor cancer patient prognosis^2^. The mechanisms underlying CIN remain poorly characterized but reflect abnormalities in kinetochore-microtubule attachment, sister chromatids cohesion, centrosome duplication, telomeres or the spindle assembly checkpoint (SAC)^3^.

To maintain genome integrity, the cell cycle must be tightly coordinated to ensure the faithful transmission of hereditary information between generations. Cells spend more than 90% of their time in interphase and interphase length correlates with total length of the cell cycle^4^. In contrast, mitosis is extremely short, and the time spent in mitosis is remarkably constant and uncoupled from variability in other phases. Mitosis is the process by which a cell properly divides its genetic material and consists of five active phases from prophase to telophase. The major mitotic checkpoint comprises the metaphase-to-anaphase transition, separating mitotic entry and exit. Coordination of the mitotic regulatory network relies on hierarchal phosphorylation cascades driven by cyclin-dependent kinases (CDK)^5,6^ and the ubiquitin-proteasome system (UPS) under the control of the anaphase-promoting complex (APC/C)^7,8^. During mitosis, known degradative functions are mainly restricted to UPS, specialized in ubiquitin-triggered protein degradation and presumably faster than lysosome-dependent degradation.

Lysosomes are acidic cytosolic vesicles responsible to enzymatically degrade all types of biological material. During macroautophagy, double-membrane vesicles (autophagosomes) engulf cytosolic material, converge and fuse with lysosomes. The lysosomal proton pump v-ATPase drives lumen acidification while the BORC-associated protein complex, including KIF5B motor protein, is the main driver of anterograde lysosomal transport and also contributes to autophagosome-lysosome fusion^9–11^. Autophagy-dependent degradation of cargos can either be non-selective (bulk) or targeted (selective). Adaptor proteins, such as p62 (also known as Sequestosome-1) regulate selective autophagic degradation^12,13^. Being cytosolic vesicles, research on autophagy and lysosomes has mainly focused on interphase cells, omitting their implication on mitosis. As cells undergo mitosis, dramatic structural rearrangements of organelles occur^14–16^. However, several recent studies show controversial observations about the function of lysosomes and autophagy in mitosis. Some studies claim that autophagy signaling is shut-down at mitotic entry, that autophagic structures are barely detected in dividing cells and that proteasome-dependent degradation of WIPI2 upon mitotic induction suppress autophagic flux^17–19^. In contrast, lysosomal-dependent degradation is implicated in Cyclin A2 proteolysis during mitosis^20^ and mitophagy is active in prophase^21,22^. ULK1, an autophagy initiator protein, was shown to drive SAC recruitment to kinetochores through Mad1 phosphorylation^23^. Furthermore, selective autophagic degradation of centriolar satellites was recently demonstrated to support correct karyokinesis^24,25^. Loss of Beclin1, another key autophagic protein, induced gene amplification and consequent aneuploidy^26^. Better understanding of the regulatory mechanisms driving mitotic transitions is crucial for preventing CIN and developing novel cancer treatment strategies.

Here, we show, for the first time, that correct mitotic progression does not rely only on UPS-dependent degradation. Dissecting the mitotic transition, we define a novel function of selective autophagy and lysosome-dependent degradation specifically in dividing cells and identify new mitotic lysosomal substrates. Furthermore, impairment of lysosome function during cell division triggers chromosome mis-segregation and induces a striking nuclear phenotype, the toroidal nucleus, which provides a new tool for genotoxicity tests.

## RESULTS

### Lysosomes and autophagic vesicles are present and active during cell division

To investigate the presence of lysosomes in mitotic subphases, LAMP2-positive vesicles and DAPI-stained DNA were analyzed by immunofluorescence. Here we showed that lysosomes are present and dynamic in all mitotic subphases (**Fig.1A; Video 1**). Morphological analysis of mitotic lysosomes indicated that lysosomes decreased in number while increased in size from prophase to late telophase, when they started recovering their interphase-like morphology (**Fig.1A-C; Video 1**). In addition, lysosome distribution is modified once cells proceeded into cell division. Mitotic lysosomes surrounded the chromosomes in prophase and moved to the edges of chromosomes during anaphase/telophase until chromosomes decondensed (**Fig.1D**). To analyze whether mitotic lysosomes maintain their degradation capacity during cell division, we stained live U2OS cells stably expressing Histone 2B-GFP (H2B-GFP U2OS) with Lysosensor for acidic organelle detection and MagicRed for lysosomal cathepsin B activation. Colocalization between Lysosensor and MagicRed supported the presence of functional lysosomes in dividing cells (**Fig.1E**).

**Figure 1.**
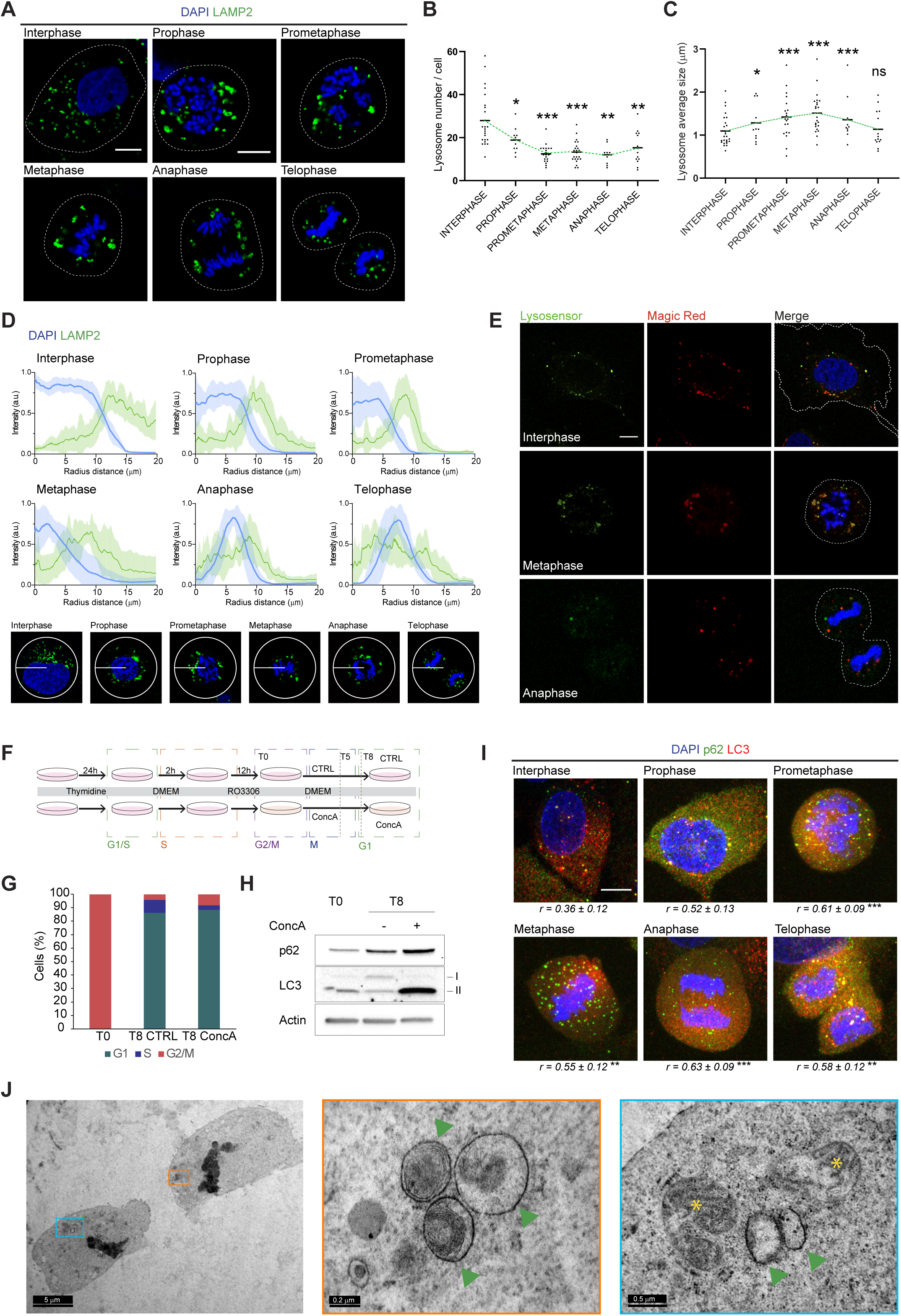
Autophagic flux and lysosome-dependent degradation are active in dividing cells. (**A**) Representative single focal plan (1Z) confocal images of U2OS cells undergoing mitosis labelled with lysosomal marker LAMP2 (green). Interphase cell and distinct mitotic subphases are detectable with DNA staining with DAPI (blue). Scale bar, 10 µm. (**B**) Quantification of lysosome number per cell in interphase compared to each mitotic subphases. Error bars represent S.D. of ≥ 10 images. (**C**) Quantification of lysosome average size per cell in interphase and for each mitotic subphases. Error bars represent S.D. of ≥ 10 images. (**D**) Upper panel: Analysis of the distribution of lysosomes (green) and DNA (blue) in interphase and for each mitotic subphases using Radial Profile Angle ImageJ plugin. A Radius of 20 μm was maintained constant for the analysis. Lower panel: representative image of a cell in each analyzed phase. (**E**) Representative single focal plan confocal images of U2OS H2B-GFP stable cells in interphase or undergoing mitosis were stained with Lysotracker (green – arbitrary color) for lysosomes detection, Magic Red (red) for active cathepsin B, H2B-GFP (blue – arbitrary color) for DNA staining. Scale bar, 10 µm. (**F**) Schematic representation of the synchronization protocol established for U2OS cells. (**G**) Cell cycle analysis of cells at T0 (before release of the reversible CDK1 inhibitor – RO3306) and of cells after release from RO3306 incubated either with normal growing media (CTRL) or with 10 nM ConcA-containing growing media. Percentage of cells in G1, S and G2/M cell cycle phases are represented under the experimental conditions. (**H**) Analysis of autophagic flux by Western Blot detection with specific antibodies of autophagic proteins p62 and LC3 in cell fractions of panel G synchronized as in panel F. β-actin protein level was used as loading control. (**I**) Representative maximal projection (z ≥ 15) of confocal images of U2OS cells in interphase or cells undergoing mitosis. Endogenous autophagic proteins LC3 (red) and p62 (green) were detected by immunofluorescence and DNA was marked with DAPI (blue). Pearson’s correlation coefficient (r) was calculated for each analyzed condition (n ≥ 10). Scale bar, 10 µm. (**J**) TEM images show mitotic cells undergoing mitotic exit. Arrowheads indicate autophagic vesicles and lysosomes and yellow asterisks point out mitochondria. Scale bars, as indicated. **Panels B** and **C**, statistical significance between interphase and mitotic subphases is represented as: * p < 0.05, ** p < 0.005, *** p < 0.001

As we found that lysosomes are present and active in mitosis, we analyzed the autophagic flux during cell division. To this end, the expression of p62 and the lipidated form of LC3 (LC3-II) were analyzed in mitosis after inhibition of lysosomal acidification with ConcA. Established synchronization protocols such as serum starvation or double-thymidine block were insufficient to efficiently synchronize U2OS cells in G2. Therefore, based on the ability of CDK1-specific inhibitor RO3306 to reversibly block cells in G2, we established a synchronization-release protocol to obtain enriched mitotic population (**Fig.1F**). Cell cycle analysis validated the efficiency of the protocol in control cells or upon lysosome inhibition (**Fig.1G**). Upon ConcA treatment, synchronized cells showed specific accumulation of LC3-II and of the autophagic adaptor protein p62 without major changes in the global cell cycle profile (**Fig.1G-H**). Time-course assay after cell synchronization and release demonstrated that both autophagic proteins LC3-II and p62 gradually accumulated during mitosis transition upon acidification blockade (**Fig.S1A-B**). In parallel, immunofluorescence analysis of both endogenous LC3 and p62 corroborated the presence of autophagic vesicles in dividing cells (**Fig.1I**). Indeed, Pearson’s correlation coefficient (Rr) between LC3 and p62 significantly increased during mitotic subphases compared to interphase cells (**Fig. 1I**). Finally, double-membrane vesicles (autophagosomes), as well as dense single-membrane vesicles (autolysosomes/lysosomes), were detected by Transmission Electron Microscopy (TEM) in mitotic cells (**Fig.1J**).

Collectively, our results demonstrate that both autophagic vesicles and functional lysosomes are present and active in mitotic cells.

### Lysosome acidification capacity and trafficking maintain correct mitotic progression

To investigate the role of lysosomes in cell division, we studied mitotic cells with impaired lysosomes either by inhibiting their degradative capacity or their intracellular trafficking. Impairment of lysosome acidification by the v-ATPase inhibitor Concanamycin A (ConcA) led to a reduced number of mitotic lysosomes while increasing lysosome size, according to their defective degradation capacity (**Fig.2A, C-D**). Parallel to ConcA treatment, we assessed the effect of KIF5B depletion on lysosome morphology, number and distribution in mitotic cells. While in interphase, KIF5B depletion induced a dramatic clustering of lysosomes in the perinuclear region and a reduction in the lysosomal number. However, these effects were negligible in mitotic cells. Interestingly, KIF5B depletion did not affect either lysosome number or size in dividing cells (**Fig.2B-E and Fig.S2A**). When we analyzed the lysosomal distribution in the different mitotic subphases, we observed that the hierarchical distribution of lysosomes observed in control cells was almost completely absent in KIF5B-depleted cells (**Fig.2E**). Interestingly, in this setting, KIF5B did not induce a clustering of lysosomes in a central zone as in interphase but impeded the spatial organization of lysosomes during mitosis suggesting the implication of this kinesin on the heterogeneous distribution of lysosomes during mitosis (**Fig.2E**).

**Figure 2.**
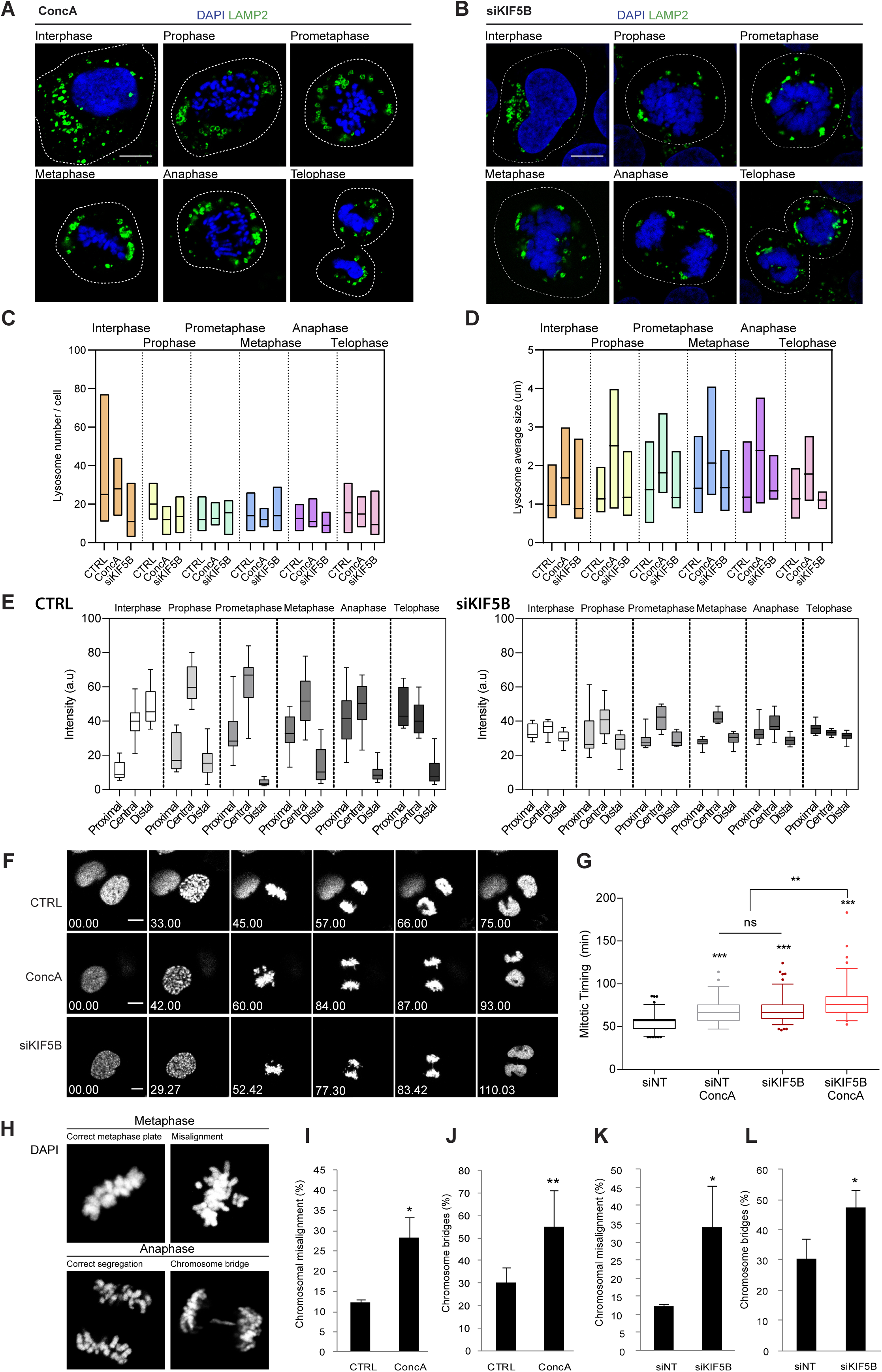
Lysosome impairment delays mitotic progression and leads to mitotic errors and an atypical nuclear phenotype. (**A**-**B**) U2OS cells were either treated with ConcA 10 nM for 24 hours (**A**) or transiently transfected with silencing RNA against KIF5B (siKIF5B) (**B**). Single plan confocal images show LAMP2-positive lysosomes (green) detected by immunofluorescence in cells undergoing mitosis. DNA is labelled with DAPI (blue). Scale bar, 10 µm. (**C**-**D**) Comparison of the number of lysosomes per cell (**C**) or the average lysosome size (**D**) in interphase and each mitotic subphases in control, ConcA-treated or KIF5B-depleted cells. Line represents the mean in each condition. Total images analyzed ≥ 10 per condition. (**E**) Intracellular distribution of lysosomes in interphase and in each mitotic subphases in control and KIF5B-depleted cells. Total images analyzed ≥ 10 per condition. (**F**) H2B-GFP U2OS cells were subjected to time-lapse imaging for 24 hours every 6 minutes 33 seconds. Single focal plan of representative images of control cells, cells treated with ConcA 10 nM or depleted for KIF5B (siKIF5B) for 48 hours undergoing mitosis are shown. H2B-GFP is depicted in grayscale (arbitrary color). Scale bars, 5 µm. (**G**) Quantification of mitotic timing of the indicated experimental conditions was performed from prophase (chromosomes condensation) to telophase (chromosomes decondensation). Error bars represent 5-95 percentiles of mitosis ≥ 90. (**H**) Representative images of misaligned metaphase plate and chromosome bridges. Scale bar, 5 µm. (**I-J**) U2OS cells were synchronized at late G2 phase and released with or without ConcA for 1 hour (**I**) or 3 hours (**J**). DAPI staining was used for DNA detection. Chromosome misalignment (**I**) and chromosomal bridges (**J**) were quantified compared to total number of metaphases and anaphases, respectively. Error bars represent S.D. of n=3 independent experiments (>200 cells). (**K**-**L**) U2OS cells depleted for KIF5B (siKIF5B) or transfected with siRNA control (siNT) were synchronized and released for 1 hour (**K**) or 3 hours (**L**) to determine the percentage of chromosome misalignment or chromosome bridges relative to the total number of cells undergoing metaphase or anaphase, respectively. Error bars represent S.D. of n=3 experiments (>150 cells). In **Panels G** and **I-L**, statistical significance is represented as: * p < 0.05, ** p < 0.005, *** p < 0.001.

Mitotic timing is strictly regulated and remarkably constant among different cell types and species^4^, taking approximately one hour in normal mitosis. Thus, after identifying active lysosomes in mitotic cells the direct implication of lysosomes in mitotic progression was tested. To this end, we used live-imaging to analyze H2B-GFP U2OS cells treated with ConcA or KIF5B-depleted and recorded mitotic timing from prophase (chromosome condensation) to telophase (chromosome decondensation). Mitosis was 22% slower in ConcA-treated cells compared to control, and KIF5B depletion delayed mitotic progression to the same extent (**Fig.2F-G; Videos 2-4**). Combination of ConcA-treatment with KIF5B knockdown increased the average mitotic duration by 45% verse control (**Fig.2G**).

In all, integrity of both lysosome transport and functionality is key for preserving mitotic schedule. To precisely characterize the involvement of lysosomes in mitotic progression, we separately analyzed the timing of mitotic entry (prophase to metaphase) and exit (metaphase to telophase). Interestingly, both episodes were delayed by treatments that impaired lysosomes (**Fig.S2B**). Consistently, the additive effect was reflected in both mitotic entry and exit (**Fig.S2B**).

One of the main causes of mitotic delay is the accumulation of mitotic errors such as misaligned chromosomes at metaphase plate or chromosome bridges appearing during chromosome segregation in anaphase^27^ (**Fig.2H**). Thus, we next quantified the acquirement of mitotic errors in synchronized cells treated or not with ConcA. Our results demonstrated that lysosome acidification blockade significantly induces the accumulation of misaligned chromosomes and chromosome bridges (**Fig.2I**). In parallel, we tested whether, beyond acidification, the protective role of lysosomes against mitotic errors also relies on trafficking. Consistent with the observed mitotic delay (**Fig.2F-G**), KIF5B-depleted synchronized cells showed a higher frequency of both types of mitotic errors (**Fig.2J**). Thus, active lysosomes are important organelles to ensure correct mitosis progression and limit CIN signature.

### Mitotic errors induced by lysosome impairment correlate with an abnormal nuclear phenotype: the toroidal nucleus

Defective mitotic progression generated a robust and striking nuclear phenotype in interphase cells. Precisely, we observed a DAPI-stained nucleus with a hole devoid of chromatin (**Fig.3A**). To understand how this structure was originated, we followed the change on nuclear structure during mitosis by live-imaging using H2B-GFP U2OS cells. Our results showed that this unconventional nuclear phenotype forms upon mitosis, being detectable in at least one of the daughter cells once chromosomes decondense (**Fig.3B; Video 5**). To assure that this unconventional structure does not correspond to an enlarged nucleolus^28^, immunofluorescence analysis was performed and results showed that none of the tested nucleolar markers (nucleolin, fibrillarin and UBF) co-stained with the DAPI-free area (**Fig.3C** and **S3A**). To resolve whether this section is nuclear or cytosolic, we imaged the nuclear envelope by immunofluorescence using Lamin B1 and Nuclear Pore Complex (NPC) antibodies (**Fig.3D** and **S3B**). Interestingly, the nuclear envelope was correctly formed around DNA. Further immunofluorescence analysis showed that cytoskeleton filaments such as Phalloidin-stained actin fibers and microtubules, as well as LAMP2-positive lysosomes or LC3-p62-positive autophagosomes were present within e void at the center of the nuclei (**Fig.3E** and **S3C-D**). To discard a nuclear invagination and/or an artifact, 3D reconstruction of high-quality images using IMARIS software resulted in the characterization of a donut-like shaped structure that we refer to as a “toroidal nucleus” (**Fig.3F** and **Video 6**). Ultrastructure analysis of these toroidal nuclei by TEM confirmed that cytosolic material, including autophagic vesicles and lysosomes were present within the hole (**Fig.3G**).

**Figure 3.**
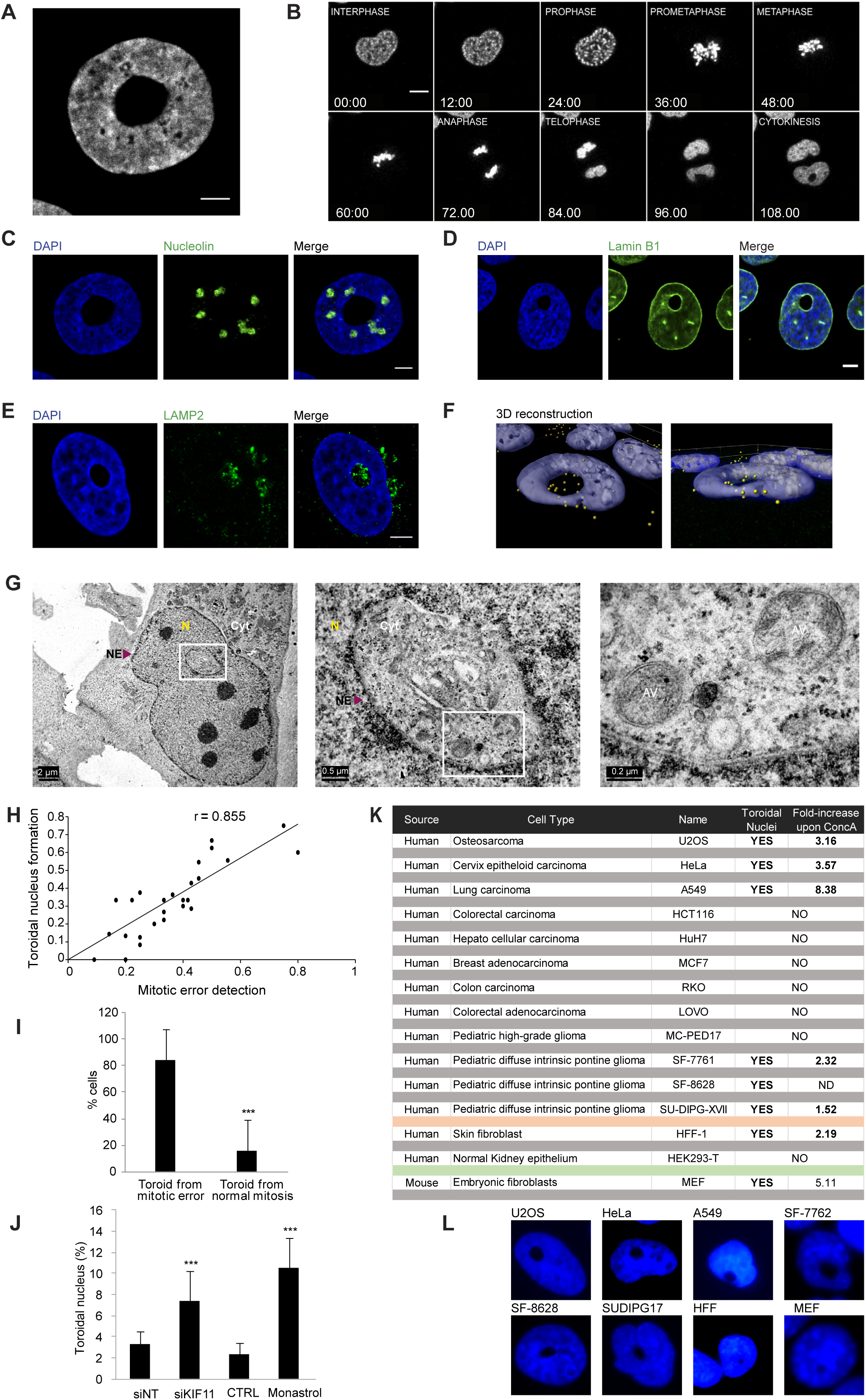
Toroidal nucleus is a novel biomarker for chromosomal instability in interphase cells. (**A**) Representative image of toroidal nucleus. U2OS cells were fixed and stained with DAPI for DNA detection. Scale bar, 5 µm. (**B**) Live imaging of H2B-GFP U2OS cells undergoing cell cycle for 16 hours every 93 seconds. Mosaic of single focal plan images indicate aberrant nucleus formation upon cell division. Scale bar, 10 µm. (**C**-**E**) U2OS cells were fixed and immunofluorescence was performed using nucleolin antibody for nucleolus detection (**C**), Lamin B1 antibody for nuclear envelope staining (**D**) or LAMP2 antibody to mark lysosomes (**E**) and DAPI for DNA labelling. Scale bar, 5 µm. (**F**) 3D reconstruction using IMARIS software was performed from confocal images stack of toroidal nucleus (z every 0.1 µm for 11 µm). Blue corresponds to DAPI-stained nucleus and yellow dots mark LAMP2-positive lysosomes. (**G**) U2OS cells were treated with ConcA for 24 hours, fixed and prepared for TEM. Images show toroidal nucleus surrounded by the nuclear envelope. Abbreviations used: NE: nuclear envelop, N: nucleus, Cyt: cytosol, AV: autophagic vesicles. Scale bars, as indicated. (**H**) Analysis of live imaging experiment by following cell division of H2B-GFP U2OS cells for 24 hours every 9 minutes. Mitotic cells were detected and visualized to quantify the number of those with detectable mitotic errors (misaligned chromosomes or chromosomal bridges) and those generating toroidal nucleus. Linear regression was analyzed and the corresponding Pearson coefficient (r) was calculated. (**I**) Analysis of the percentage of toroidal nucleus formed upon non-apparent mitotic error or as a consequence of mitotic errors. Error bars represent S.D. of 295 mitosis. (**J**) U2OS cells were depleted for KIF11 or MCAK for 48 hours using silencing RNA or treated with Monastrol 100 µM or Nocodazole 1 µM for 24 hours. Quantification of toroidal nuclei formed under these conditions was performed by detection of DAPI-stained nuclei. Error bars represent S.D. of n > 3 experiments (10 fields / experiment). (**K**) Table summarizing a screen for toroidal nuclei through various cell lines. Fold-increase was calculated upon ConcA treatment (10 nm for 24 h). (**L**) Representative images of toroidal nuclei in screened cell lines from panel K. Panels **I** and **J** statistical significance is represented as: * p < 0.05, ** p < 0.005, ***p < 0.001.

This phenotype was once reported as donut-shaped nucleus^29^. The authors proposed that protein farnesylation inhibitors (FTI) induce the formation of toroidal nuclei by causing a pericentrin-related centrosome separation defect in HeLa cells. Thus, we tested whether FTI induced the formation of toroidal nuclei in our model. After confirming the production of toroidal nuclei following FTI treatment in HeLa cells (**Fig.S3E, left**), we demonstrated that U2OS cells did not respond to FTI. This is contrast to our results with impairment of lysosome acidification by ConcA, which significantly increased the percentage of toroidal nuclei in both cell types (**Fig.S3E, right**). To verify that FTI does not stimulate toroidal nuclei formation with a different kinetics in U2OS, we treated and analyzed cells for 24h, 48h and 72h with FTI or ConcA. While ConcA gradually induced a robust increase of toroidal nuclei with time; FTI did not (**Fig.S3F**). Next, we analyzed whether inhibition of lysosome acidification by ConcA or Chloroquine (CQ) induced the inhibition of farnesylation of Lamin A/C. Neither ConcA nor CQ do not impede Lamin A/C farnesylation like FTI (**Fig.S3G**). Our results suggest that the formation of toroidal nuclei is not due to the inhibition of farnesylation *per se* but to a defect in mitotic progression regardless its inducer.

Mitotic stresses have been linked to prolonged mitosis and accumulation of mitotic errors, both of which we detected after lysosomal impairment. As toroidal nuclei were formed following karyokinesis, we examined by live imaging a putative link between the occurrence of mitotic errors and the formation of toroidal nuclei. The results showed a linear correlation between detectable mitotic errors and toroidal nucleus generation with a Pearson’s correlation coefficient of 0.85 (**Fig.3H**). Indeed, single-cell analysis demonstrated that more than 80% of toroidal nuclei resulted from cells undergoing apparent defective mitosis (**Fig.3I**). Based on this, we next investigated the role of mitotic stresses on generating toroidal nuclei. To this end, we depleted cells of KIF11, a kinesin required for bipolar spindle establishment^30^, and analyzed the formation of toroidal nuclei. Disruption of KIF11 by small interfering RNA (siKIF11) or by the KIF11 specific inhibitor Monastrol induced a significant increase in toroidal nucleus frequency (**Fig.3J**) without dramatically affecting lysosome morphology and localization (**Fig.S3H**). Depletion of other kinesins implicated in chromosome segregation such as mitotic centromere-associated kinesin (siMCAK/KIF2C)^31^ or Centromere-associated protein E (siCENP-E)^32^ produced a 3.3- and 3.8-fold increase of toroidal nucleus population, respectively (**Fig.S3I**). In all tested conditions, lysosomes morphology or distribution were not apparently perturbed (**Fig.S3J**). Furthermore, cells treated with Nocodazole, a microtubule de-polymerizing agent, showed a 4.5-fold increased prevalence of toroidal nuclei accompanied by the expected lysosomal collapse (**Fig.S3I-J**). Thus, the formation of toroidal nucleus seems to be attributable to impairment of karyokinesis.

We next investigated whether this phenotype is a consequence of impaired nuclear envelope reformation. Live imaging of H2B-GFP U2OS cells stably expressing mCherry-Lamin A/C, indicated that the reformation of the nuclear envelope preceded the formation of toroidal nuclei (**Fig.S3K; Video 7**). This nuclear phenotype was versatile in terms of size, nuclear localization and morphology (**Fig.S3L1-4**). Furthermore, the nucleus could contain more than one void (**Fig.S3L5-8**) and be accompanied by micronucleus (**Fig.S3L6**). After mitosis, one or both daughter cells could harbor this phenotype (**Fig.S3L7-8**).

We investigated whether toroidal nucleus was a common feature or specific to U2OS cells. To this end, we screened through a panel of cell lines. Toroidal nuclei were not detected under the tested conditions in the colon carcinoma cell lines RKO, LoVo or HCT116, hepatocarcinoma cells Huh7 or MC-PED17 glioma-derived cell line^33–36^ (**Fig.3K**). However, detection of toroidal nuclei was successful in various cells, among which non-transformed human skin fibroblast (HFF) and embryonic mouse fibroblasts (MEF), as well as cells from lung carcinoma (A549), cervix carcinoma (HeLa) or diffuse intrinsic pontine glioma (SU-DIPG-XVII, SF-8628 and SF-7761)^33,34,37^ (**Fig.3K-L**). All those cells, except SF-8628, responded to ConcA treatment by presenting a significant increase of toroidal nuclei (**Fig.3K**).

Thus, toroidal nuclei can be scored as a read-out for mitotic errors in interphase cells.

### Lysosome disruption induces the formation of toroidal nuclei

The presence of toroidal nuclei in interphase cells facilitates the analysis of mitotic impairment in whole cell populations, favoring the toroidal nucleus as a powerful tool for quantitative analysis of chromosomal instability. Here we aimed to screen for lysosome-specific stresses using toroidal nucleus as a biomarker for chromosomal instability. To this end, cells were treated with ConcA or depleted for KIF5B and toroidal nuclei frequency was quantified. Consistently, v-ATPase inhibition as well as blockage of anterograde transport led to a robust increase of toroidal nuclei population (**Fig.4A-B**).

**Figure 4.**
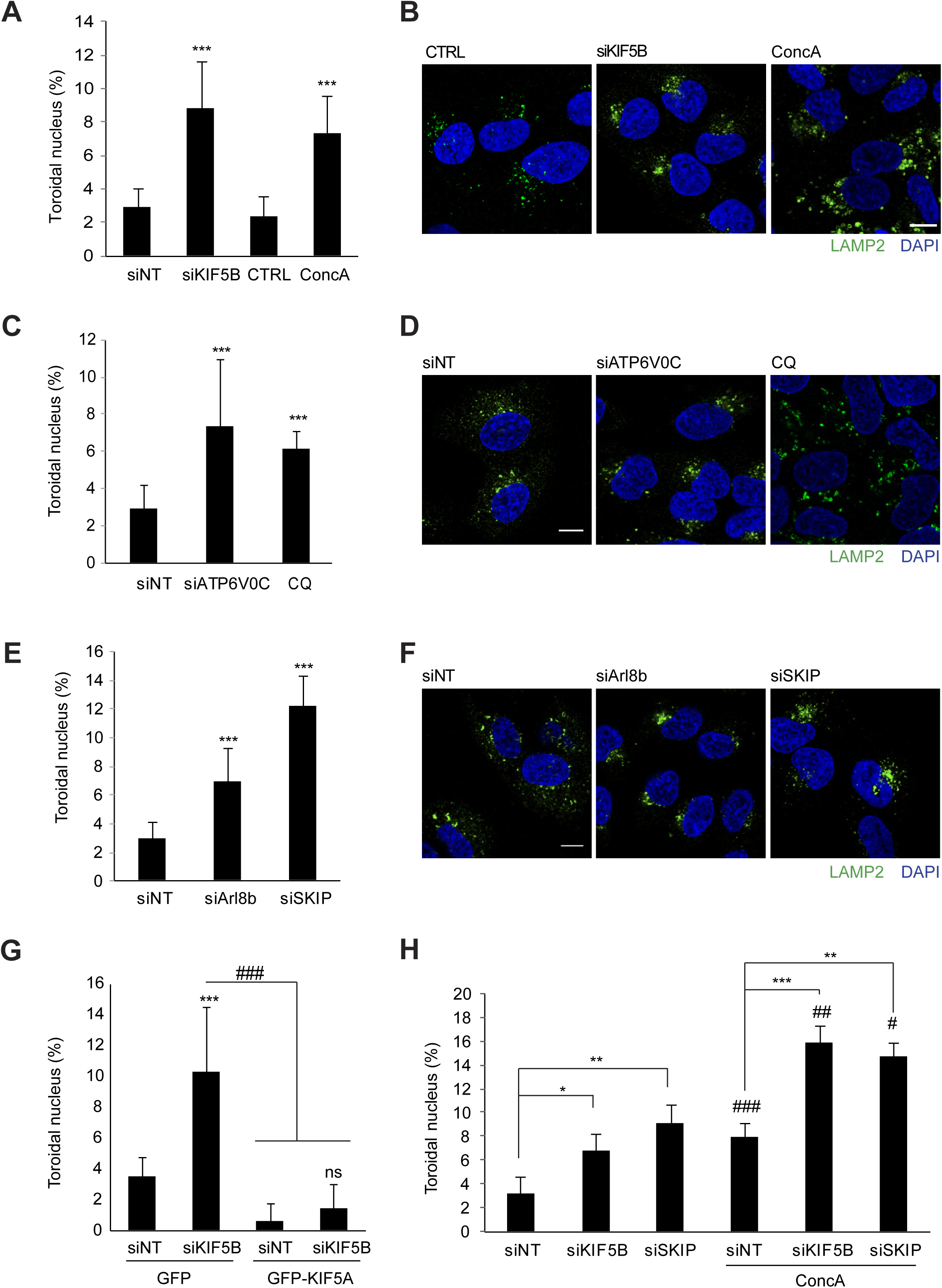
Toroidal nucleus frequency increases upon impairment of lysosome trafficking and acidification capacity. (**A**) U2OS cells were treated with ConcA (10 nM) for 24 hours or depleted for KIF5B for 48 hours using silencing RNA. Error bars represent S.D. of n > 5 experiments. (**B**) Representative confocal single plans showing morphology and distribution of endogenous lysosomes (LAMP2 in green) under experimental conditions of panel A. Nuclei were labelled with DAPi in blue. Scale bar, 10 µm. (**C**) U2OS cells were treated with Chloroquine CQ (10 µM) or depleted for the V0c subunit of the v-ATPase (siATP6V0c) for 48 hours. Error bars represent S.D. of n > 3 experiments. (**D**) Representative single plan confocal images of U2OS cells treated as in panel **C**. Lysosomes were detected with LAMP2 antibody and nuclei were stained with DAPI. Scale bar, 10 µm. (**E**) U2OS cells were depleted for BORC-associated proteins SKIP or Arl8b using silencing RNA for 48 hours. Error bars represent S.D. of n > 3 experiments. (**F**) Representative images of LAMP2-positive lysosomes (green) under the indicated conditions (panel E). Nuclei are stained with DAPI (blue). Scale bar, 10 µm. (**G**) U2OS cells were transfected with KIF5B silencing RNA and/or with KIF5A-GFP overexpression plasmid. Error bars represent S.D. of n > 3 experiments. (**H**) U2OS cells were depleted for KIF5B or SKIP and the next day treated or not with ConcA for 24 hours. Error bars represent S.D of n > 3 experiments. In panels **A, C, E** and **G-H** nuclei were stained with DAPI for detection of toroidal nuclei compared to total number of cells. Statistical significance is represented as: * p < 0.05, ** p < 0.005, ***p < 0.001, ns: p > 0.05.

To discard ConcA-side effects not related to v-ATPase inhibition, we genetically or chemically inhibited lysosome acidification. Thus, cells were treated with Chloroquine or depleted of the V0c v-ATPase subunit (siATP6V0c). In agreement, inhibition of lysosome acidification increased the formation of toroidal nucleus by 2.5-3-fold (**Fig.4C**). To validate treatment efficiency, lysosome positioning and size were assessed by immunofluorescence, confirming the expected increase in lysosomal volume^38^ (**Fig.4D**). Both ConcA and Chloroquine increased toroidal nucleus formation but their kinetics differed (**Fig.S4A**). Indeed, ConcA treatment acted faster and produced a peak effect at 24 hours, while Chloroquine produced a similar effect after 48 hours, inducing the maximum impact at 72 hours (**Fig.S4A**). To corroborate the specificity of mitosis impairment to lysosome-related stresses, we tested compounds targeting other cellular processes or organelles. In contrast to lysosomal inhibition, treatment with the mitochondrial complex I inhibitor Phenformin or the proteasome inhibitor MG132 had no significant effect on the frequency of toroidal nuclei in unsynchronized cells (**Fig.S4B**). We further asked whether defects in BORC-dependent lysosome trafficking would also induce formation of toroidal nuclei as we determined in cells depleted for KIF5B (**Fig.4E**). Cells genetically depleted of BORC-associated proteins Arl8b or SKIP also showed a significant increase in the population of toroidal nuclei (2.3- and 4.1-fold respectively) confirming the importance of lysosomal trafficking for correct mitotic progression (**Fig.4F**). Depletion efficacy was corroborated by the perinuclear clustering of lysosomes (**Fig.4F**). Additionally, we examined the specificity of the motor function of KIF5B in the formation of toroidal nucleus by complementing control or KIF5B-depleted cells with neuronal KIF5A, which resulted in a rescue of normal toroidal nucleus frequency (**Fig.4G**). Interestingly, depletion of other motor proteins involved in lysosomal trafficking such as KIF3A, KIF2A or KIF1A also led to a significant increase in toroidal nuclei (**Fig.S4C**). Parallel to a delay in mitotic timing (**Fig.2F-G**), we found that simultaneous disruption of lysosome acidification (ConcA) and trafficking (siKIF5B or siSKIP) had an additive effect on toroidal nucleus formation compared to single treatments (**Fig.4H**). In all, our results confirm that both v-ATPase-dependent lysosome acidification and BORC-associated lysosomal trafficking are required to preserve mitotic integrity.

Impairment of lysosome functionality does not prevent mitotic cells to progress into G1 phase but leads to CIN. In cells synchronized and released specifically in mitosis, ConcA and Chloroquine similarly led to toroidal nuclei formation (2.6- and 2.7-fold increase, respectively) (**Fig.S4D**), indicating that the different kinetics observed might be due to effect of the drugs in other phases of the cell cycle. Interestingly, neither MG132 nor Monastrol induced toroidal nuclei in these conditions (**Fig.S4D**), but it is noteworthy that these drugs imped cells from properly concluding cell division as previously demonstrated^39,40^, thus limiting the generation of subsequent daughter cells harboring toroidal nucleus (**Fig.S4E**). These results further suggest that lysosomes are not required to undergo mitosis, but genomic instability is triggered when lysosomes are impaired.

### Macroautophagy is a key player to maintain mitosis fidelity

As we have detected the presence of autophagic vesicles in mitosis (**Fig.1**), we next analyzed the involvement of macroautophagy for faithful mitotic progression by scoring for toroidal nucleus frequency after macroautophagy inhibition. Genetic depletion of Atg5 (siAtg5) induced a significant increase in the proportion of toroidal nuclei (1.6-fold increase) (**Fig.5A**). Notably, ConcA did not significantly affect the formation of toroidal nucleus in Atg5-depleted cells (**Fig.5A**). In agreement with the correlation between mitotic errors occurrence and formation of toroidal nuclei, the inhibition of autophagy by Atg5 depletion provoked a significant increase in misaligned chromosomes and chromosome bridges (**Fig.5B-C**). Protein depletion efficiency was monitored by Western Blot (**Fig.S5A**). Chemical inhibition of autophagy by 3MA, a PI3K-class III inhibitor, also resulted in an increase in toroidal nuclei frequency together with an enhanced detection of chromosomal misalignment (**Fig.S5B-C**). As U2OS is a cancer cell line already susceptible to genomic instability and chromosome alterations^41^, we used mouse embryonic fibroblasts (MEFs) to analyze the impact of defective autophagy on mitotic progression in a non-tumoral model. First, we validated ConcA efficiency and autophagic flux impairment by analyzing Atg5 and LC3 protein levels. As expected, Atg5-/-MEFs harbored a defective autophagic flux (**Fig.5D**). Then, we quantified the frequency of toroidal nuclei in Atg5 wt (Atg5+/+) and Atg5 KO (Atg5-/-) MEFs. ConcA treatment in Atg5+/+ MEFs significantly increased toroidal nuclei population, demonstrating that lysosomal inhibition impacts mitotic progression in various cell lines (**Fig.5E-F and Fig.3K-L**). Like the effect of the depletion of Atg5 in U2OS cells, Atg5-/-MEFs presented a significantly increased percentage of toroidal nuclei under normal growing conditions, supporting that basal autophagy prevents cells from mitotic errors. ConcA-induced toroidal nucleus formation was not further increased in Atg5 KO MEFs (**Fig.5E**). Atg5-deficient MEFs were significantly more susceptible to chromosomal misalignment and chromosomal bridges compared to parental MEFs (2.36- and 1.48-fold increase respectively) (**Fig.5G-H**). Inhibition of v-ATPase-dependent lysosomal acidification by ConcA clearly increased chromosome misalignment and chromosomal bridges in control Atg5+/+ MEFs (**Fig.5G-H**). Cells lacking Atg5 showed a milder response to inhibition of lysosomal acidification regarding both phenotypes (**Fig.5G-H**).

Altogether, our results establish the involvement of macroautophagy machinery in correct mitotic progression.

**Figure 5.**
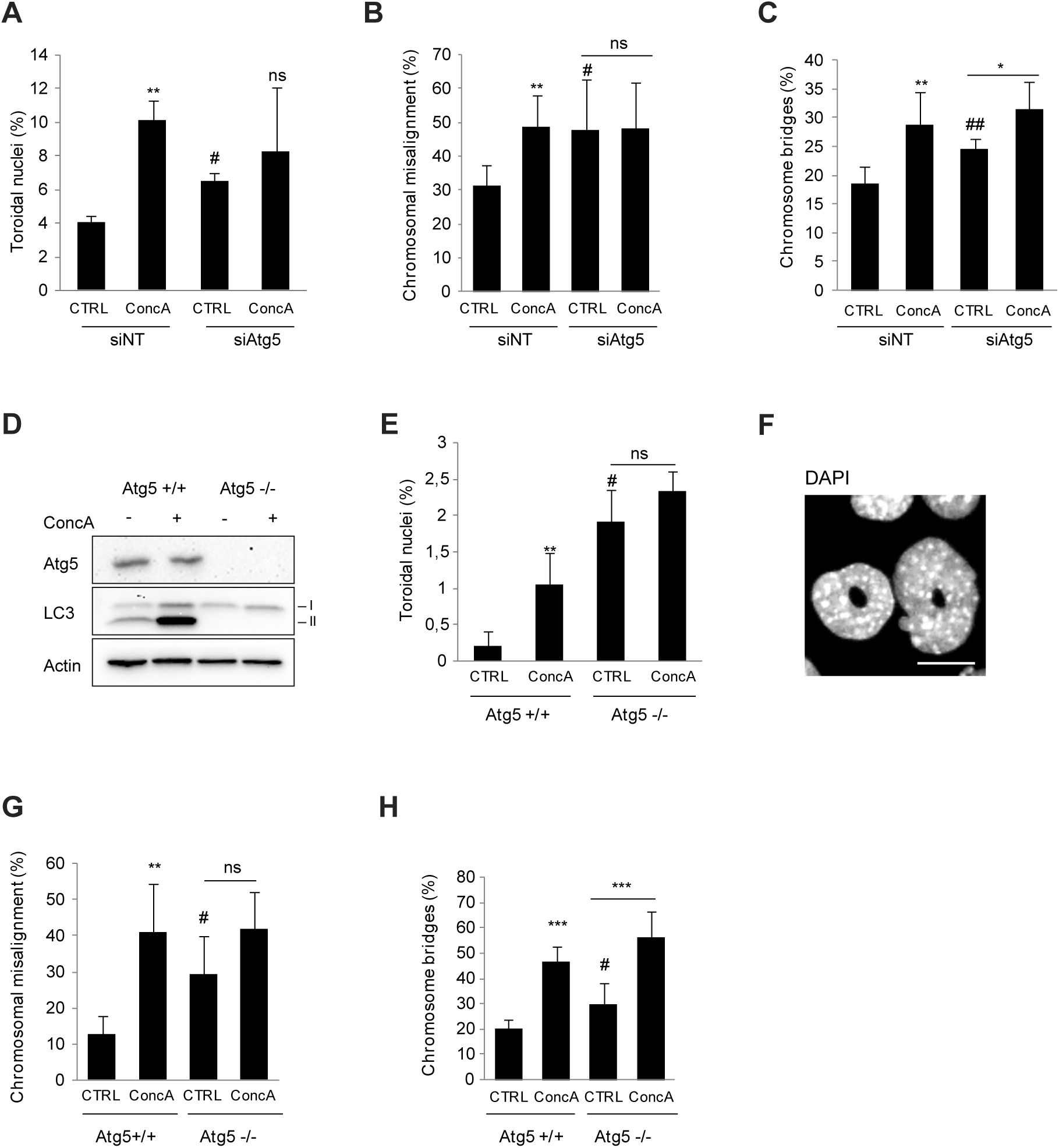
Alterations of macroautophagy increase mitotic errors and toroidal nucleus formation. (**A**) U2OS cells were depleted for Atg5 for 48 hours and treated or not with ConcA for 24 hours. Error bars represent S.D. of n > 3 experiments. (**B-C**) Quantification of mitotic errors (chromosome misalignment and chromosome bridges, respectively) in U2OS cells control (siNT) or depleted for Atg5 (siAtg5) for 48 hours under normal growing condition (CTRL) or treated with ConcA (10 nM for 24 h). Error bars represent S.D. of n=4 experiments (>300 cells). (**D**) Validation of Atg5 depletion and ConcA efficacy in MEFs Atg5 +/+and -/- by Western Blot. β-actin was used as loading control. (**E**) MEF Atg5 +/+ and Atg5 -/- were subjected or not to ConcA treatment for 24 hours. DNA was labelled with DAPI and frequency of toroidal nuclei was quantified. Error bars represent S.D. of n=3 experiments. (**F**) Representative image of toroidal nucleus in MEF Atg5 KO cells. Scale bar, 5 µm. (**G-H**) Quantification of mitotic errors (chromosome misalignment and chromosome bridges, respectively) in MEF Atg5 +/+ and Atg5 -/-. Error bars represent S.D. of n=5 experiments (>100 cells). Panels **A**-**C, E** and **G**-**H**, * represents the statistical significance of ConcA inhibitory effect compared to control, while # represents the effects of Atg5 genetic depletion. * p < 0.05, ** p < 0.005, *** p < 0.001, ns: p > 0.05, # p < 0.05 and ## p < 0.005.

### Cohesin proteins WAPL and PDS5B are novel lysosome substrates during cell division

Based on the characterization of a remarkable role for lysosomes and autophagic vesicles in mitotic progression, we next aimed to identify novel substrates of the autophagic-lysosomal pathway specifically in mitosis by mass spectrometry analysis. Following synchronization of cells at G2/M transition, cells were released in normal media or in media containing ConcA for 7 hours. G1-enriched cell fractions were processed for mass spectrometry analysis. Drug efficacy was validated by LC3 and p62 accumulation in the three experimental replicates after ConcA treatment (**Fig.S6A**). Proteomic data analysis identified a total of 1749 peptides present in both ConcA and control fractions, while only 13 and 141 were uniquely detected in control and ConcA fractions, respectively (**Fig.6A**). Enrichment analysis validated 853 peptides being differently expressed (q-value<0.05) that were selected based on ± 1.2-fold change cut-off (**Fig.6B**). In concordance with its role in inhibiting lysosome-dependent degradation, ConcA significantly induces protein accumulation. The obtained list of candidates was clustered into cellular and organism functions using Reactome free database^42,43^. Reactome analysis highlighted that proteins accumulated after lysosome inhibition specifically during mitosis were mainly involved in cell cycle, vesicle-mediated transport or DNA repair (**Fig.S6B**). In agreement with previous data (**Fig.1I, S1B** and **S6A**), p62 was significantly increased by 2.73-fold in the ConcA fraction compared to control (**Fig.6B**). Next, we selected the 56 protein candidates that were clustered in functional annotations related to mitosis and cell cycle regulation and we performed a GO enrichment analysis (**Fig.S6C**). A list of 20 proteins involved in mitotic cell cycle progress (GO:1903047) were further studied to define the involvement of lysosomes in chromosome segregation (**Fig.6C**). Interestingly, among the principal enriched proteins, two cofactors of the cohesin complex such as Cohesin-associated factor B (PDS5B) and Wings apart-like protein (WAPL) were significantly increased by 4.78- and 1.64-fold, respectively (**Fig.6B-C**). To validate proteomic data, we isolated mitotic cells by shake-off after synchronization-release and evaluated protein levels in control and ConcA-treated cells. As expected, autophagic proteins p62 and LC3-II accumulated in ConcA-treated mitotic fraction (**Fig.6D and S6D**). Significant increase of both WAPL and PDS5B protein levels occurred in mitotic cells upon acute impairment of lysosomal acidification capacity (**Fig.6D-E**), indicating that both PDS5B and WAPL are novel putative lysosomal substrates specifically during mitotic progression. To validate our findings in another cell line, we obtained shake-off fractions from non-transformed MEFs. Mitotic fractions of non-synchronized MEFs corroborated that PDS5B and WAPL significantly accumulated more in Atg5 -/- MEFs cells compared to parental MEFs (**Fig.6F-G and S6E**). WAPL and PDS5B accumulation upon lysosome inhibition could potentially be due to the observed mitotic delay independent of the autophagy machinery. To address this point, U2OS cells were transfected with exogenous WAPL-GFP protein and treated or not with ConcA for 3 hours after synchronization. Mitosis-enriched fractions were then subjected to GFP-immunoprecipitation. First, we validated the binding between WAPL and its well-described partner PDS5B^44^ (**Fig.6H**). WAPL-GFP interacts with PDS5B to the same extent in control or ConcA-treated cells, suggesting that their interaction is maintained upon lysosomal inhibition. Interestingly, WAPL-GFP interaction with autophagic adaptor protein p62 significantly increased in ConcA-treated cells (**Fig.6H**). These results support that WAPL is recognized by p62-dependent selective autophagy machinery and emerges as a novel lysosome substrate during mitosis.

**Figure 6.**
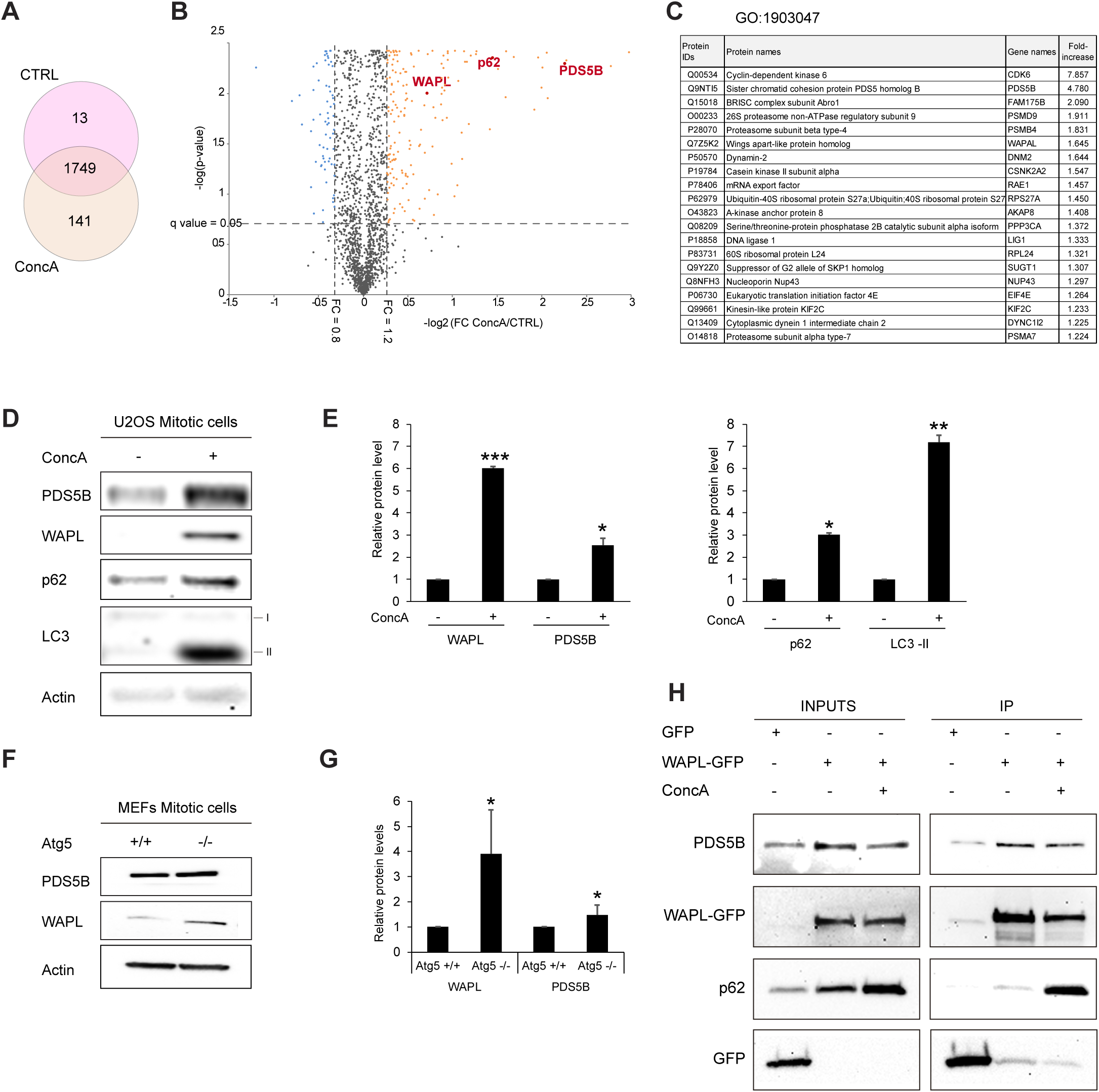
Identification of novel mitotic lysosomal substrates. (**A**) Venn diagram representing proteins detected by mass spectrometry of synchronized U2OS cells released in growing media (CTRL) or ConcA-containing media. (**B**) Volcano plot depicting significant fold changes (FC) (log_2_) in protein abundances in cells treated with 10 nM ConcA. Three independent experiments were analyzed per condition. Dots denote the 1749 unique proteins detected in pre-filtered samples. Threshold at q-value < 0.05 depicts peptides significantly modified under experimental conditions. Blue dots represent peptides with FC < 0.8. Orange dots correspond to peptides presenting a FC > 1.2. Within these candidates, red dots indicate target proteins WAPL, p62 and PDS5B. (**C**) Table summarizing the 20 candidates identified by mass spectrometry that are clustered in the Gene Ontology (GO) group GO:1903047 of mitotic cell cycle progress. (**D**) Western Blot of synchronized U2OS mitotic cell fractions after mitotic cell enrichment by shake-off. Protein levels of WAPL, PDS5B, p62 and LC3 were detected. β actin was used as loading control. (**E**) Quantification of WAPL, PDS5B, p62 and LC3-II protein levels in experimental conditions as in panel D and normalized to β actin protein level. Error bars represent S.D. of n=3 experiments. (**F**) MEFs Atg5 +/+ and -/- under normal growing conditions were subjected to shake-Off. Mitotic enriched cell fractions were then analyzed by Western-Blot and accumulation of WAPL and PDS5B was detected in MEF Atg5 -/-. β-actin was used as loading control. (**G**) Quantification of WAPL and PDS5B protein levels from panel F normalized to β-actin loading control. Error bars represent S.D. of n=3 experiments. (**H**) Immunoprecipitation of exogenous WAPL-GFP in U2OS cells treated or not with 10 nM ConcA for 24 hours. Direct interaction between WAPL and p62 was detected by Western-Blot. Cells expressing exogenous GFP tag were used as negative control. Statistical significance is represented as: * p < 0.05, ** p < 0.005, ***p < 0.001, ns: p > 0.05.

In all, our results shed light into a novel regulatory mechanism for accurate mitotic progression, depicting lysosomes as key players for correct chromosomal congression and faithful genetic transmission (**Fig.7**). In addition, we identified the toroidal nuclei as a read-out of mitotic defects in interphase cells, and such, are a novel biomarker of genotoxicity for cancer research.

**Figure 7.**
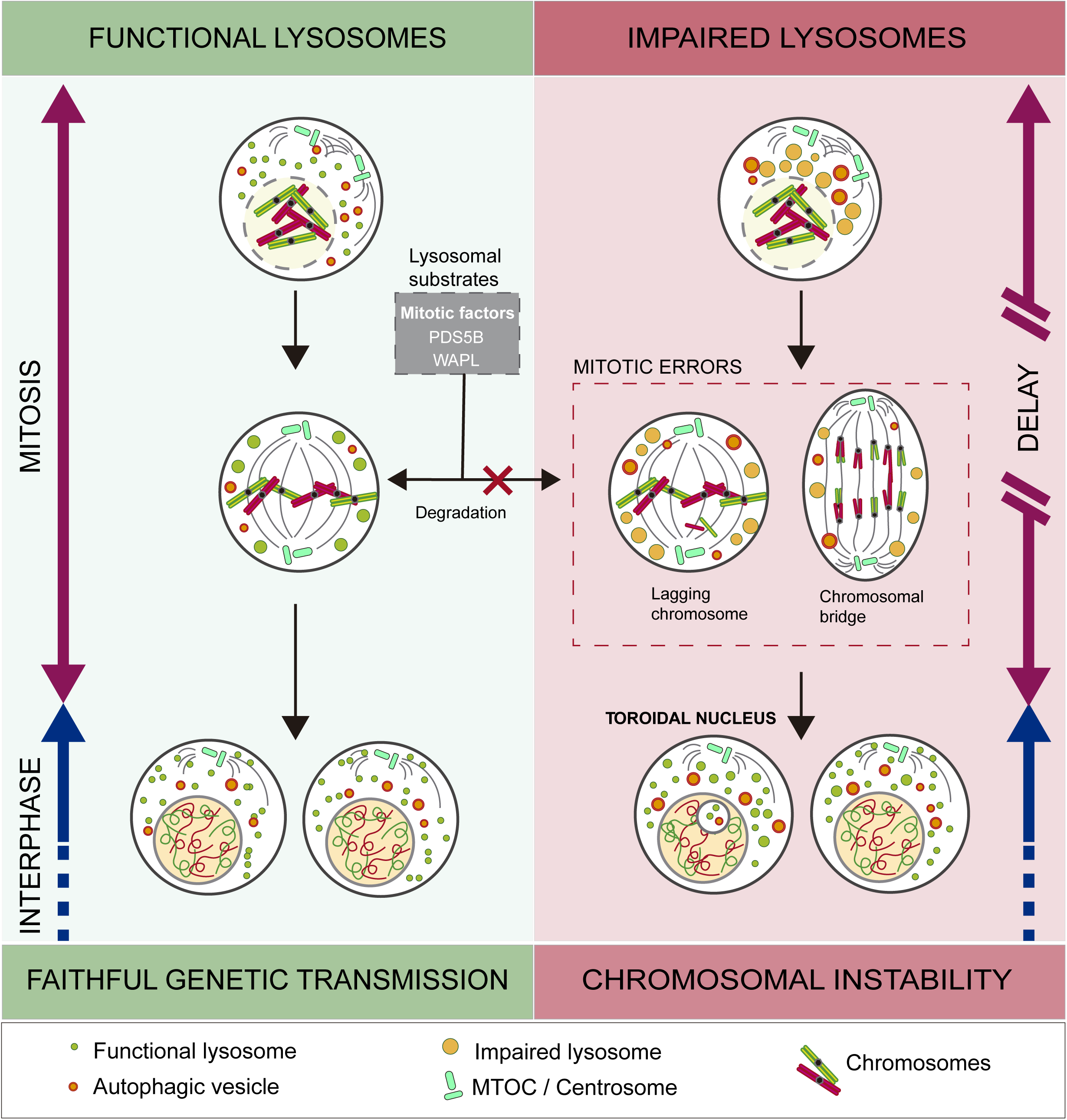
Impairment of lysosomes in mitotic cells leads to chromosomal instability. Lysosomal function is required for the maintenance of correct cell division and faithful transmission of genetic material in the two daughter cells. Functional lysosomes degrade essential mitotic factors necessary for appropriate chromosome segregation. On the contrary, when lysosomes are dysfunctional either by impairment of their trafficking or acidification capacity, mitotic cells are prompt to accumulate mitotic errors, to generate toroidal nucleus and mitosis is delayed. These alterations are key features of chromosomal instability.

## DISCUSSION

### Faithful mitotic progression relies on autophagy and lysosomes

During mitosis, dramatic cellular rearrangement occurs to support proper chromosome segregation and productive partitioning of intracellular organelles. Many membranous compartments undergo massive spatiotemporal disruption^16^ and scheduled protein degradation is required. While the role of UPS has been extensively studied, the implication of lysosomes and autophagy in cell division still remains elusive^17,18,20–23,45^. Although lysosome studies are mainly focused on interphase cells, endocytic vesicles (including lysosomes) were proposed to serve as a membrane source for plasma membrane extension and retraction during mitotic transition^14,15^. Recently, PCM1-driven selective degradation of centriolar satellites was shown to maintain centrosome function for correct mitotic progression^24^. Here, our results agree with the persistence of a robust autophagic flux and lysosomal-dependent degradation during mitosis. Lysosome enlargement suggests that fusion events are occurring in early mitosis and that, in contrast to other organelles, lysosomes play an active role during cell division. The finding that under basal conditions Atg5 KO MEFs harbor a higher population of toroidal nuclei corroborates that autophagy is active and necessary for mitosis even in non-transformed cells. Among the identified lysosomal substrates in mitosis, the accumulation of p62, an autophagic adaptor protein for ubiquitinated cargos, validates the persistence of autophagic flux and suggests a potential alternative route for the degradation of mitotic factors^46,47^. To note, PCM1 was not detected in our proteomic analysis of mitotic cells, supporting that various selective autophagy processes might probably coexist and should be timely regulated during mitotic progression. Recently, p62 emerged as a potential pivotal player in the crosstalk between UPS and selective autophagosome-lysosome degradation^48,49^. Based on p62 bifunctionality and ubiquitin-lysine linkages diversity, we speculate a potential interconnection between the two main degradative pathways to correctly distribute genetic material. Control of sister chromatid cohesion during cell division is crucial for equal distribution of genetic material. The canonical evolutionary conserved cohesin complex is one of the main molecular entities involved in this process. The cohesin complex consists of four subunits (SMC1, SMC3, SCC1 and STAG1/3)^50^. Currently, most research focuses on cohesin loading processes during replication and cohesin-dependent extrusion of chromatin loops for DNA-damage repair^51^. In mitosis, cohesin supports both mitotic entry and metaphase-to-anaphase transition. In prophase, cohesin complexes dissociate to allow chromatid condensation and separation. At that precise moment, cohesin complex disruption is triggered by WAPL and PDS5B along the chromatids arms^52,53^. WAPL-dependent dissociation at the centromeric region is inhibited by Shugoshin to prevent premature chromatin separation^54,55^. During metaphase-to-anaphase transition, chromosome dissociation at the centromeric region is regulated by APC/C-dependent securin ubiquitination, leading to separase activation that cleaves SCC1 to culminate chromatids separation^56^. Since lysosomes can degrade most biological material and, taking in account that in this study we identified more than 100 potential lysosomal protein substrates in mitotic cells, we favor the idea that the implication of lysosomes in cell division is a multifactorial process. Our finding that WAPL and PDS5B accumulate upon blockade of lysosomal acidification, identifies them as potential lysosomal substrates specifically during mitosis. Our study reports for the first time the involvement of autophagy- and lysosomal-dependent degradation of cohesin cofactors. The identification of WAPL as a novel p62 interactor supports the model of selective degradation of WAPL by lysosome during mitotic transition. However, additional studies would be needed to better understand the interconnection between lysosome-dependent degradation and the regulation of chromatid cohesion during mitotic progression. Variations in PDS5B and WAPL expression correlate cohesin and chromosomal segregation defects with aneuploidies and cancer progression^52,57–60^.

While the implications of lysosome biology in cancer are still controversial, efficacy of lysosomotropic agents has been proven in several malignancies either alone or in combination, and clinical trials are ongoing^61,62^. CIN is considered a hallmark of cancer understood either as a strength, due to mutation induction facilitating tumor diversification, or as a tumor vulnerability by eventually triggering apoptotic cell death^2,63^. Defective autophagy was recently linked to increased CIN^64^. Here we show that lysosomal acidification and trafficking as well as selective autophagy are necessary to prevent CIN. Our data are consistent with the metaphor of autophagy and lysosome-dependent degradation as a double-edged sword in cancer progression^65^. In all, depending on cell type and tumor stage, a combination of lysosomotropic drugs with conventional therapies might be beneficial, as autophagy inhibition can be detrimental for cancer cells and increased CIN can lead to cell death. Therefore, better understanding of the mechanisms by which selective autophagy functions in cell division will provide novel routes to improve cancer treatment strategies.

### Toroidal nucleus, a new tool for genotoxicity screenings

Study of mitosis is challenging due to the velocity and architectural changes accompanying cell division. To escape these obstacles, most mitosis-focused studies take advantage of synchronization protocols. However, drugs used for cell synchronization, such as Nocodazole, do not specifically target mitosis but rather essential cellular components like microtubules, thus altering basic cellular functions. Currently, the only marker of defective mitosis detectable in non-mitotic cells is the micronucleus. Although micronuclei formation is a consequence of chromosome missegregation^66^, they can be reabsorbed during subsequent cell divisions or destroyed by autophagy^67,68^. Therefore, complementary read-outs in interphase cells are needed for the analysis of mitosis impairment. Here we show that toroidal nuclei form upon inexact mitosis and linearly correlates with the occurrence of mitotic errors. Inhibition of farnesylation was previously proposed to drive the formation of donut-shaped nuclei by affecting centrosome function^29^. Here we demonstrate that toroidal nuclei do not specifically and uniquely form upon inhibition of protein farnesylation. Indeed, FTI did not increase toroidal nucleus population in U2OS, while mitotic stresses, such as inhibition of lysosomal function described here, significantly and robustly increased the frequency of toroidal nuclei in various cell lines. In addition, our results reveal that toroidal nuclei form prior to reformation of the nuclear envelope, as it was also described for micronucleus^69^. The fate of micronuclei is still uncertain. Four possibilities are under examination: degradation, reincorporation into the nucleus, extrusion or persistence in the cytoplasm leading to apoptosis or chromothripsis^70^. We expect similarities for toroidal nuclei, but further analyses are needed to decipher how cells can cope with toroidal nuclei. We detected toroidal nuclei in transformed and non-transformed cells, designating presence of this structure as a robust and conserved phenotype within cell lines. However, the occurrence of toroidal nuclei might vary dependent on cell type specificities and/or genetic background. Variability in toroidal nucleus frequency might also correlate with basal autophagy levels, which are known to be cell type specific^71^. Noteworthy, detection of toroidal nuclei is a new convenient tool for impaired mitosis in genotoxicity screenings.

Until now, orchestration of mitotic factors degradation has been exclusively attributed to the UPS. Our data reveals an additional route in which active autophagy and lysosomes promote faithful genetic transmission. Given the promising results of lysosomotropic drugs for cancer treatment, our study opens an alternative line of research focusing on lysosomes modulation to exploit CIN for cancer therapy.

## MATERIALS AND METHODS

### Cell Culture

U2OS, HeLa, A549, HCT116, RKO, Huh7, MCF7 and HEK293-T cell lines were obtained from American Type Culture Collection (ATCC). Atg5 wt and Atg5 KO MEF were kindly provided by Dr. Zorzano (IRB Barcelona). HFF cells were kindly provided by the ICO CellBank. U2OS H2B-GFP cell line was kindly provided by Dr. Agell (UB). SF-7761 and SF-8628 were purchased from Millipore Sigma (Burlington, MA). MC-PED17 cells were a gift from Dave Daniels (Mayo Clinic; Rochester, MN). SU-DIPG-XVII cells were a gift from Michelle Monje (Stanford University; Palo Alto, CA). U2OS mCherry-LaminA/C and U2OS H2B-GFP LAMP1-RFP were generated in our laboratory by transient transfection of plasmids purchased in Addgene repository, LAMP1-GFP (#34831), LAMP1-mRFP-Flag (#34611) and mCherry-LaminA/C (#55068). WAPL-GFP plasmid was kindly shared by Dr. Fangwei Wang. Briefly, DNA transfection was performed following manufacturer’s instructions using Lipofectamine 2000 (Life Technologies) in 1:5 opti-MEM: DMEM medium. Cells were selected using a fluorescence-activated cell sorter (MoFlo Atrios SORTER). Cells were grown in DMEM high glucose (Gibco) [4mM L-Glutamine, 4.5 g/L glucose, 1 mM Pyruvate] supplemented with 10% heat-inactivated fetal bovine serum (FBS) (Sigma Aldrich). Cells were incubated at 37°C, 5% CO2 and 90-95% of relative humidity. Specific experimental conditions are indicated in figure legends. All chemicals used in this study are reported in **Table 1**.

### Cell synchronization at G2

Cells were seeded the day before starting the synchronization in DMEM medium supplemented with FBS (complete DMEM). Thymidine was added at a final concentration of 2 μM for 24 hours to synchronize cells at late G1. After PBS washing, cells were released to S phase by adding complete DMEM for 2 hours. Finally, RO3306 (CDK1 inhibitor) was added to cell culture medium at a final concentration of 9 μM for 12 hours to arrest cells at late G2 (**Fig.1F**). Experiments with synchronized cells were performed by releasing cells in complete DMEM with or without the corresponding drugs for the indicated times. For the obtention of shake-off fractions, synchronized cells were subjected to consecutive strokes to the dishes to detach specifically the mitotic cells while maintaining the integrity of the cell monolayer.

### Silencing RNA transfections

siRNA transfections were performed following manufacturer’s instructions in Opti-MEM medium (Life Technologies) using Lipofectamine RNA-iMAX (Life Technologies). Unless otherwise indicated, transfections were performed for 48 hours. siRNA sequences and concentrations used in these studies are listed in **Table 2**.

### Cell Cycle analysis

Cells were trypsinized, counted and placed on ice. 500,000 cells were centrifuged at 1200 rpm at 4°C for 5 minutes and cell pellet was washed with ice-cold PBS, fixed with 70% ethanol and placed at −20°C for at least 24 hours. Cells were then washed with ice-cold FACS buffer [BSA 0.1%, EDTA 5 mM in PBS] and centrifuged for 5 minutes at 1000 rpm at 4°C. Supernatant was discarded and the cell pellet was resuspended in propidium iodide (PI) staining solution [PBS, 0.1% NP40, 20 µg/mL RNAse A (Invitrogen), 40 µg/mL PI (Sigma)]. Cell suspension was transferred to a 5 mL tube with cell strainer cap (Corning) and maintained at room temperature protected from light for 15 minutes. Samples were acquired using FACS Canto System (BD Biosciences, USA) and analyzed with ModFit LT software (Verity Software House).

### Protein extraction and Western Blot

Cells were washed twice with ice-cold PBS, scraped and lysed on Lysis Buffer [20 mM Tris-HCl pH 8, 10 mM EDTA, 150 mM NaCl, 1% Triton-X100] supplemented with protease inhibitor cocktail (1:100 Sigma-Aldrich) and phosphatase inhibitors cocktail 2 and 3 (1:100 Sigma-Aldrich). Cell lysates were centrifuged at 13000 rpm for 10 minutes at 4°C. Protein concentration was analyzed using Pierce BCA Protein Assay kit (ThermoFisher Scientific) following manufacturer’s instructions. Equal amounts of protein lysates were resuspended in Laemnli SDS-sample buffer and incubated at 95°C for 5 minutes. Proteins were separated on SDS-PAGE and transferred to PVDF membranes (Millipore). Membranes were blocked with 5% non-fat dry milk (BioRad) in Tris-buffered saline containing 0.1% Tween-20 (TBS-T) for 1 hour at room temperature. Incubation of primary antibodies was performed overnight at 4°C in 5% non-fat dry milk or 3.5% BSA (Sigma-Aldrich). After three washes in TBS-T, membranes were incubated for 1 hour at room temperature with secondary antibodies (1:5000) diluted in 5% non-fat milk. Upon incubation, membranes were washed three times with TBS-T and protein detection was performed by using enhanced chemiluminescence kit (GE Healthcare). Blots were scanned with iBright detection system (Thermo Fisher). All the antibodies used in this study are reported with the corresponding working concentrations in **Table 3**.

### Immunoprecipitation

U2OS cells transfected with WAPL-GFP expression vector or GFP-empty vector as negative control were washed twice with ice-cold PBS and cellular pellets were kept at −80°C before lysis. Cellular pellet was lysed in ice-cold IP-RIPA buffer [100 mM Tris-HCl pH7.5, 300 mM KCl, 10 mM MgCl_2_, 2 mM EGTA, 20% Glycerol and 1.6 % NP40] supplemented with protease inhibitors (2X), phosphatase inhibitors cocktails (1X), PMSF 1mM and DTT 1mM. After 5 minutes incubation on ice, collected samples were mechanically lysed with cold syringe and centrifuged at 13,000 rpm for 5 minutes at 4 °C. Soluble fraction was subjected to BCA Pierce protein quantification. 1,500 μg of protein per sample were separated for immunoprecipitation at 1 μg/μl concentration. For the immunoprecipitation, 30 μl of GFP-Trap beads (ChromoTek) were added to each lysate and incubated by rotation overnight at 4 °C. Immunoprecipitates were washed three times with IP-RIPA buffer. Immunoprecipitated proteins were denatured by the addition of 2X sample buffer followed by boiling for 10 minutes, resolved by 4 %-20 % Criterion TGX Gel (BIO-RAD) electrophoresis and analyzed by immunoblotting. 20 μg of the total lysate were loaded as input to control.

### Immunofluorescence

Cells were grown as monolayers on coverslips and subjected to the indicated conditions. Cells were fixed with 4% paraformaldehyde for 10 minutes at room temperature. After 5 minutes washing with PBS, cells were permeabilized using 0.1% Triton X-100 in PBS for 10 minutes at room temperature. Next, cells were blocked with 1% BSA/0.01% Triton X-100 in PBS plus 10 mM Glycine for 30 minutes at room temperature. Cells were incubated with primary antibodies for one hour at room temperature or overnight at 4°C. Following a series of PBS washes, cells were incubated with secondary antibodies for 45 minutes at room temperature. After two washes of 5 minutes with PBS, coverslips were mounted using Vectashield Mounting Solution containing DAPI (Vector Laboratories). All the antibodies used in this study are reported with the corresponding working concentrations in **Table 2**.

### Vesicle acidification and detection assays in live cells

U2OS cells were seeded onto glass coverslips and subjected to the indicated conditions. Live cells were washed once with PBS and Magic Red (Immunochemistry Technologies) and Lysosensor (Invitrogen) were added to cover the cell layer following manufacturer’s instructions. Cells were incubated for 10 minutes at room temperature before image acquisition.

### Image acquisition and analysis

For detailed analysis, image acquisition was performed in Leica spectral confocal microscope TCS SP5 using a 63x N.A 1.4 objective and LAS AF software or Carl Zeiss LSM880 confocal microscope and ZEN software. Fluorophores were excited with Argon 488, DPSS 561, Diode 640 and Diode 405 lasers. Image analysis was performed using FIJI Image J software (NIH USA).

For toroidal nucleus and mitotic errors quantification, images were acquired using Nikon Epifluorescence microscope using a 40x dry objective. Image analysis was performed using Cell counter plugin from FIJI Image J software (NIH USA). Pearson’s correlation coefficient was calculated based on confocal images using FIJI ImageJ plugin Intensity Correlation Analysis. Mitotic lysosomes size and number was analyzed with FIJI ImageJ fixing particles size from 0.3 μm^2^ to infinity. The distribution of mitotic lysosomes was calculated with FIJI ImageJ plugin Radial Profile Angle as previously described^72^.

### 3D reconstitution

Stacks of images acquired at optimal settings with Carl Zeiss LSM880 confocal microscope were processed with IMARIS Software (Bitplane) for 3D image reconstruction and to generate in silico animations and images.

### Live-cell time-lapse videos

For mitotic delay analysis, U2OS H2B-GFP cells were grown onto glass bottom 8-well slides (IBIDI). After indicated treatments, medium was replaced by FluoroBright medium (Gibco) and live-cell imaging was performed on the Leica spectral confocal microscope TCS SP5. Images were taken every 6 minutes and 33 seconds for a total time of 24 hours using the 63x glycerol objective.

For lysosome trafficking analysis, U2OS cells stably expressing LAMP1-RFP and H2B-GFP were grown onto glass bottom 8-well slides (IBIDI). After indicated treatments, medium was replaced by FluoroBrite medium (Gibco) and live-cell imaging was performed on Carl Zeiss LSM880 confocal microscope. Images were taken every 5 minutes for a total time of 24 hours using the 63x glycerol objective.

For toroidal nucleus and nuclear lamina reformation assays, U2OS cells stably expressing H2B-GFP and mCherry-Lamin A/C were grown onto bottom-glass 8 chambers slides (IBIDI). After addition of FluoroBrite with indicated treatments, mitotic cell division was analyzed on a Carl Zeiss LSM880 confocal microscope. For toroidal nucleus formation, images were taken every 93 seconds for 16 hours using the 63x glycerol objective. For nuclear envelope reformation experiment, images were taken every 5 minutes for 16 hours using the 63x glycerol objective.

### Transmission Electron Microscopy (TEM)

U2OS cells with or without ConcA (10 nM) treatment for 24 hours were fixed with Glutaraldehyde 2.5% in Sodium Cacodylate Trihydrate 0.1 M pH 7.2 for 2 hours at 4°C. After three washes with Sodium Cacodylate Trihydrate 0.1 M of 15 minutes each, samples were incubated for 2 hours with Osmium tetroxide 1% in Sodium Cacodylate Trihydrate 0.1 M at room temperature. Samples were washed three times for 15 minutes with Sodium Cacodylate Trihydrate 0.1 M. Then, samples were processed through dehydration with Ethanol 30% to 100% gradually. Samples were then embedded into Resin EPOXY. After sample orientation, polymerization occurred at 60°C for 48 hours. Samples were cut using ultramicrotome EM UC6 Leica, first in semithin of 250 nm and then ultrathin of 70 nm. Image acquisition was performed with TEM microscope JEOL JEM-1011. Images were analyzed and processed with FIJI ImageJ software (NIH).

### Mass Spectrometry

Three replicates of treated samples were processed for protein extraction in RIPA lysis buffer [Tris-HCl pH8 50mM, NaCl 150mM, NP-40 1%, Sodium deoxycholate 0.5%, SDS 0.1%]. Protein concentration was determined and 50 μg of each sample was digested using a FASP (Filter-Aided Sample Preparation) approach. Briefly, proteins were reduced with dithiothreitol 10 mM (60 minutes, 32°C) and alkylated with iodoacetamide 20 mM (30 minutes at 25°C in the dark). Samples were loaded onto an Amicon Ultra filter 10 KDa, 0.5 mL (Millipore) to remove interfering agents with 2 rounds of centrifugations/washes with 100 mM ammonium bicarbonate buffer (13,600 g; 25 minutes at room temperature). Digestion was carried out in two steps: first, samples were digested (1:50 w sample/w enzyme) with Lys-C (Wako) in 6 M urea buffer for 3 hours at 35°C, second, the samples were diluted 10-fold with 100 mM ammonium bicarbonate buffer and digested with modified porcine trypsin (Promega-Gold) (1:25 w sample/w enzyme) for 16 hours at 37°C. The resulting peptide mixture was recovered by centrifuging the filter. Then, the filter was washed twice with 300 μL of 50 mM ammonium bicarbonate and once with 200 μL of 20% acetonitrile/50 mM ammonium bicarbonate (13,600g for 25 min at room temperature). All the fractions were pooled, and the final peptide mixture was acidified with formic acid. Finally, the final volume of the acidified peptide solution was reduced on a SpeedVac vacuum system (Thermo Fisher Scientific), and the peptide solution was desalinated with a C18 spin column (Thermo Fisher Scientific) following supplier’s indications.

Samples were analyzed in a Proxeon 1,000 liquid chromatographer coupled to an Orbitrap Fusion Lumos (Thermo Fisher Scientific) mass spectrometer. Samples were re-suspended in 0.5% formic acid in water, and 2 μL (1 μg/ μL) were injected for LC-MSMS analysis. Peptides were trapped on an NTCC-360/75-3-123 LC column and separated using a C18 reverse phase LC column-Easy Spray (Thermo Fisher Scientific). The gradient used for the elution of the peptides was 1% to 35% in 90 minutes followed by a gradient from 35% to 85% in 10 minutes with 250 nL/min flow rate. Eluted peptides were subjected to electrospray ionization in an emitter needle (PicoTipTM, New Objective, Scientific Instrument Services) with an applied voltage of 2,000 V. Peptide masses (m/z 300-1,700) were analyzed in data-dependent mode where a full scan MS was acquired on the Orbitrap with a resolution of 60,000 FWHM at 400 m/z. Up to the 10 most abundant peptides (minimum intensity of 500 counts) were selected from each MS scan and then fragmented using CID (collision-induced dissociation) in the linear ion trap using helium as collision gas with 38% normalized collision energy. The scan time settings were full MS at 250 ms and MSn at 120 ms. Generated raw data files were collected with Thermo Xcalibur (v.2.2) (Thermo Fisher Scientific). MaxQuant 1.6.1.0 Software (Department for Proteomics and Signal Transduction, Max-Planck Institute for Biochemistry) was used to search the raw data obtained in the MS analyses against a SwissProt/Uniprot human database with Andromeda Search engine (1.5.6.0). A target and decoy database were used to assess the false discovery rate (FDR). Trypsin was chosen as enzyme and a maximum of two misscleavages were allowed. Carbamidomethylation (C) was set as a fixed modification, whereas oxidation (M) and acetylation (N-terminal) were used as variable modifications. Searches were performed using a peptide tolerance of 7 ppm and a product ion tolerance of 0.5 Da. Resulting data files were filtered for FDR <1%. Statistical analysis was performed in Perseus 1.6.2.1 (Department for Proteomics and Signal Transduction, Max-Planck Institute for Biochemistry).

### Statistical Analysis

Data was analyzed by Excel or GraphPad Prism4 software. Results are presented as Mean ± S.D., for the indicated n independent experiments. Experimental data sets were compared by: (i) Two-sampled, two-tailed Student’s t-test for two experimental conditions sharing normal distribution and variance (ii) One-way ANOVA test for more than 2 conditions sharing normal distribution and variance. Multiple comparisons corrected using Bonferroni test (iii) Kruskal-Wallis test for more than 2 conditions and data without assuming normal distribution. Multiple comparisons corrected using Dunn’s test.

## Supporting information

Supplementary Figures with Legends

## ACKNOWLEDGEMENTS

We thank Dr. George Thomas, Dr. Sara Kozma and the rest of the members of the laboratory of Cancer Metabolism for scientific inputs and sharing reagents. We thank Dr. Neus Agell for providing H2B-GFP stable U2OS cells, Dr. Antonio Zorzano for MEFs Atg5 parental and KO and Dr. Cristina Muñoz for siRNAs siAtg5 and sip62. We are grateful to Dr. Juan S. Bonifacino for sharing KIF5A-GFP plasmid and to Dr. Fangwei Wang for WAPL-GFP plasmid. We also thank the technical facilities at CCITUB for FACS analysis and confocal microscopy. We are grateful to Dr. Joffrey Pelletier, Dr. Thomas Neufeld and Dr. Terje Johansen for their constructive comments on the manuscript. This study was supported by grants to AT from Ministerio de Economía, Industria y Competitividad (SAF2017-85561-R), which is part of Agencia Estatal de Investigación (Co-funded by European Regional Development Fund. ERDF, a way to build Europe), by joint grants to the Laboratory of Cancer Metabolism from Instituto de Salud Carlos III-Red Temática de Investigación Cooperativa en Cáncer (RD12/0036/0049), and from Generalitat de Catalunya-Suport als *Grups de Recerca de Catalunya* (2017SGR1743). EA was supported by Ministerio de Educación, Cultura y Deporte (FPU13/05400) and SAF2017-85561-R. C.D. was supported by Mayo Clinic/NIH training grant 5-T32-CA217836-02 and NIH grant RO1-HL125353 (through Edward Hinchcliffe). C.M. was supported by Juan de la Cierva fellowship (IJCI-2015-24716) from Ministerio de Ciencia Innovación y Universidades and by European Union’s Horizon 2020 research and innovation program under the Marie Sklodowska-Curie grant agreement (M-Lysosomes, 799000). The authors declare no competing financial interests. We thank CERCA Program/Generalitat de Catalunya for institutional support to IDIBELL.

The authors declare no conflict of interest and no competing financial interests.

## AUTHORS CONTRIBUTIONS

E.A. and C.M. conceived, designed and performed the experiments. C.D. performed experiments for detection of toroidal nuclei in glioma cells. E.A., C.M., S.A. and A.T. analyzed and discussed the data. E.A. and C.M. wrote the manuscript. A.T. and C.M. coordinated the study. All authors intellectually contributed and commented on the manuscript.

## REFERENCES

1. McClelland SE. Role of chromosomal instability in cancer progression. Endocr Relat Cancer 2017; 24:T23–31.

2. Bakhoum SF, Cantley LC. The Multifaceted Role of Chromosomal Instability in Cancer and Its Microenvironment. Cell 2018; 174:1347–60.

3. Thompson SL, Bakhoum SF, Compton DA. Mechanisms of chromosomal instability. Curr Biol CB 2010; 20:R285–295.

4. Araujo AR, Gelens L, Sheriff RSM, Santos SDM. Positive Feedback Keeps Duration of Mitosis Temporally Insulated from Upstream Cell-Cycle Events. Mol Cell 2016; 64:362–75.

5. Sullivan M, Morgan DO. Finishing mitosis, one step at a time. Nat Rev Mol Cell Biol 2007; 8:894–903.

6. Bloom J, Cross FR. Multiple levels of cyclin specificity in cell-cycle control. Nat Rev Mol Cell Biol 2007; 8:149–60.

7. Peters J-M. The anaphase promoting complex/cyclosome: a machine designed to destroy. Nat Rev Mol Cell Biol 2006; 7:644–56.

8. Pines J. Cubism and the cell cycle: the many faces of the APC/C. Nat Rev Mol Cell Biol 2011; 12:427–38.

9. Jia R, Guardia CM, Pu J, Chen Y, Bonifacino JS. BORC coordinates encounter and fusion of lysosomes with autophagosomes. Autophagy 2017; 13:1648–63.

10. Guardia CM, Farías GG, Jia R, Pu J, Bonifacino JS. BORC Functions Upstream of Kinesins 1 and 3 to Coordinate Regional Movement of Lysosomes along Different Microtubule Tracks. Cell Rep 2016; 17:1950–61.

11. Pu J, Schindler C, Jia R, Jarnik M, Backlund P, Bonifacino JS. BORC, a multisubunit complex that regulates lysosome positioning. Dev Cell 2015; 33:176–88.

12. Mancias JD, Kimmelman AC. Mechanisms of Selective Autophagy in Normal Physiology and Cancer. J Mol Biol 2016; 428:1659–80.

13. Lamark T, Svenning S, Johansen T. Regulation of selective autophagy: the p62/SQSTM1 paradigm | Essays in Biochemistry [Internet]. [cited 2019 Sep 18]; Available from: http://essays.biochemistry.org/content/61/6/609.full-text.pdf

14. Albertson R, Riggs B, Sullivan W. Membrane traffic: a driving force in cytokinesis. Trends Cell Biol 2005; 15:92–101.

15. Boucrot E, Kirchhausen T. Endosomal recycling controls plasma membrane area during mitosis. Proc Natl Acad Sci 2007; 104:7939–44.

16. Jongsma MLM, Berlin I, Neefjes J. On the move: organelle dynamics during mitosis. Trends Cell Biol 2015; 25:112–24.

17. Eskelinen E-L, Prescott AR, Cooper J, Brachmann SM, Wang L, Tang X, Backer JM, Lucocq JM. Inhibition of autophagy in mitotic animal cells. Traffic Cph Den 2002; 3:878–93.

18. Furuya T, Kim M, Lipinski M, Li J, Kim D, Lu T, Shen Y, Rameh L, Yankner B, Tsai L-H, et al. Negative regulation of Vps34 by Cdk mediated phosphorylation. Mol Cell 2010; 38:500–11.

19. Lu G, Yi J, Gubas A, Wang Y-T, Wu Y, Ren Y, Wu M, Shi Y, Ouyang C, Tan HWS, et al. Suppression of autophagy during mitosis via CUL4-RING ubiquitin ligases-mediated WIPI2 polyubiquitination and proteasomal degradation. Autophagy 2019;:1–18.

20. Loukil A, Zonca M, Rebouissou C, Baldin V, Coux O, Biard-Piechaczyk M, Blanchard J-M, Peter M. High-resolution live-cell imaging reveals novel cyclin A2 degradation foci involving autophagy. J Cell Sci 2014; 127:2145–50.

21. Li Z, Zhang X. Autophagy in mitotic animal cells. Sci Bull 2016; 61:105–7.

22. Liu L, Xie R, Nguyen S, Ye M, McKeehan WL. Robust autophagy/mitophagy persists during mitosis. Cell Cycle Georget Tex 2009; 8:1616–20.

23. Yuan F, Jin X, Li D, Song Y, Zhang N, Yang X, Wang L, Zhu W-G, Tian C, Zhao Y. ULK1 phosphorylates Mad1 to regulate spindle assembly checkpoint. Nucleic Acids Res 2019;

24. Holdgaard SG, Cianfanelli V, Pupo E, Lambrughi M, Lubas M, Nielsen JC, Eibes S, Maiani E, Harder LM, Wesch N, et al. Selective autophagy maintains centrosome integrity and accurate mitosis by turnover of centriolar satellites. Nat Commun 2019; 10:4176.

25. Joachim J, Razi M, Judith D, Wirth M, Calamita E, Encheva V, Dynlacht BD, Snijders AP, O’Reilly N, Jefferies HBJ, et al. Centriolar Satellites Control GABARAP Ubiquitination and GABARAP-Mediated Autophagy. Curr Biol 2017; 27:2123-2136.e7.

26. Mathew R, Kongara S, Beaudoin B, Karp CM, Bray K, Degenhardt K, Chen G, Jin S, White E. Autophagy suppresses tumor progression by limiting chromosomal instability. Genes Dev 2007; 21:1367–81.

27. Rieder CL, Maiato H. Stuck in Division or Passing through: What Happens When Cells Cannot Satisfy the Spindle Assembly Checkpoint. Dev Cell 2004; 7:637–51.

28. Abella N, Brun S, Calvo M, Tapia O, Weber JD, Berciano MT, Lafarga M, Bachs O, Agell N. Nucleolar Disruption Ensures Nuclear Accumulation of p21 upon DNA Damage. Traffic 2010; 11:743–55.

29. Verstraeten VLRM, Peckham LA, Olive M, Capell BC, Collins FS, Nabel EG, Young SG, Fong LG, Lammerding J. Protein farnesylation inhibitors cause donut-shaped cell nuclei attributable to a centrosome separation defect. Proc Natl Acad Sci U S A 2011; 108:4997–5002.

30. Liu Y, Zhang Z, Liang H, Zhao X, Liang L, Wang G, Yang J, Jin Y, McNutt MA, Yin Y. Protein Phosphatase 2A (PP2A) Regulates EG5 to Control Mitotic Progression. Sci Rep [Internet] 2017 [cited 2019 Mar 8]; 7. Available from: http://www.nature.com/articles/s41598-017-01915-w

31. Ishikawa K, Kamohara Y, Tanaka F, Haraguchi N, Mimori K, Inoue H, Mori M. Mitotic centromere-associated kinesin is a novel marker for prognosis and lymph node metastasis in colorectal cancer. Br J Cancer 2008; 98:1824–9.

32. Yao X, Abrieu A, Zheng Y, Sullivan KF, Cleveland DW. CENP-E forms a link between attachment of spindle microtubules to kinetochores and the mitotic checkpoint. Nat Cell Biol 2000; 2:484–91.

33. Hashizume R, Smirnov I, Liu S, Phillips JJ, Hyer J, McKnight TR, Wendland M, Prados M, Banerjee A, Nicolaides T, et al. Characterization of a diffuse intrinsic pontine glioma cell line: implications for future investigations and treatment. J Neurooncol 2012; 110:305–13.

34. Nagaraja S, Vitanza NA, Woo P, Taylor KR, Liu F, Zhang L, Li M, Meng W, Ponnuswami A, Sun W, et al. Transcriptional Dependencies in Diffuse Intrinsic Pontine Glioma. Cancer Cell 2017; 31:635-652.e6.

35. Zhang L, Peterson TE, Lu VM, Parney IF, Daniels DJ. Antitumor activity of novel pyrazole-based small molecular inhibitors of the STAT3 pathway in patient derived high grade glioma cells. PLoS ONE [Internet] 2019 [cited 2019 Sep 23]; 14. Available from: https://www.ncbi.nlm.nih.gov/pmc/articles/PMC6667205/

36. Chan K-M, Fang D, Gan H, Hashizume R, Yu C, Schroeder M, Gupta N, Mueller S, James CD, Jenkins R, et al. The histone H3.3K27M mutation in pediatric glioma reprograms H3K27 methylation and gene expression. Genes Dev 2013; 27:985–90.

37. Mueller S, Hashizume R, Yang X, Kolkowitz I, Olow AK, Phillips J, Smirnov I, Tom MW, Prados MD, James CD, et al. Targeting Wee1 for the treatment of pediatric high-grade gliomas. Neuro-Oncol 2014; 16:352–60.

38. Mauvezin C, Nagy P, Juhász G, Neufeld TP. Autophagosome–lysosome fusion is independent of V-ATPase-mediated acidification. Nat Commun 2015; 6:7007.

39. Kim OH, Lim JH, Woo KJ, Kim Y-H, Jin I-N, Han ST, Park J-W, Kwon TK. Influence of p53 and p21Waf1 expression on G2/M phase arrest of colorectal carcinoma HCT116 cells to proteasome inhibitors. Int J Oncol 2004; 24:935–41.

40. Huszar D, Theoclitou M-E, Skolnik J, Herbst R. Kinesin motor proteins as targets for cancer therapy. Cancer Metastasis Rev 2009; 28:197–208.

41. Pontén J, Saksela E. Two established in vitro cell lines from human mesenchymal tumours. Int J Cancer 1967; 2:434–47.

42. Fabregat A, Jupe S, Matthews L, Sidiropoulos K, Gillespie M, Garapati P, Haw R, Jassal B, Korninger F, May B, et al. The Reactome Pathway Knowledgebase. Nucleic Acids Res 2018; 46:D649–55.

43. Stein LD. Using the Reactome Database. Curr Protoc Bioinforma 2004; 7:p8.7.1-8.7.16.

44. Shintomi K, Hirano T. Releasing cohesin from chromosome arms in early mitosis: opposing actions of Wapl-Pds5 and Sgo1. Genes Dev 2009; 23:2224–36.

45. Pohl C, Jentsch S. Midbody ring disposal by autophagy is a post-abscission event of cytokinesis. Nat Cell Biol 2009; 11:65–70.

46. Akutsu M, Dikic I, Bremm A. Ubiquitin chain diversity at a glance. J Cell Sci 2016; 129:875–80.

47. Rogov V, Dötsch V, Johansen T, Kirkin V. Interactions between Autophagy Receptors and Ubiquitin-like Proteins Form the Molecular Basis for Selective Autophagy. Mol Cell 2014; 53:167–78.

48. Hewitt G, Carroll B, Sarallah R, Correia-Melo C, Ogrodnik M, Nelson G, Otten EG, Manni D, Antrobus R, Morgan BA, et al. SQSTM1/p62 mediates crosstalk between autophagy and the UPS in DNA repair. Autophagy 2016; 12:1917–30.

49. Nam T, Han JH, Devkota S, Lee H-W. Emerging Paradigm of Crosstalk between Autophagy and the Ubiquitin-Proteasome System. Mol Cells 2017; 40:897–905.

50. Peters J-M, Nishiyama T. Sister Chromatid Cohesion. Cold Spring Harb Perspect Biol 2012; 4:a011130.

51. Hassler M, Shaltiel IA, Haering CH. Towards a Unified Model of SMC Complex Function. Curr Biol 2018; 28:R1266–81.

52. Haarhuis JHI, Elbatsh AMO, van den Broek B, Camps D, Erkan H, Jalink K, Medema RH, Rowland BD. WAPL-mediated removal of cohesin protects against segregation errors and aneuploidy. Curr Biol CB 2013; 23:2071–7.

53. Losada A, Hirano M, Hirano T. Cohesin release is required for sister chromatid resolution, but not for condensin-mediated compaction, at the onset of mitosis. Genes Dev 2002; 16:3004–16.

54. Hara K, Zheng G, Qu Q, Liu H, Ouyang Z, Chen Z, Tomchick DR, Yu H. Structure of cohesin subcomplex pinpoints direct shugoshin–Wapl antagonism in centromeric cohesion. Nat Struct Mol Biol 2014; 21:864–70.

55. Nishiyama T, Ladurner R, Schmitz J, Kreidl E, Schleiffer A, Bhaskara V, Bando M, Shirahige K, Hyman AA, Mechtler K, et al. Sororin Mediates Sister Chromatid Cohesion by Antagonizing Wapl. Cell 2010; 143:737–49.

56. Nakajima M, Kumada K, Hatakeyama K, Noda T, Peters J-M, Hirota T. The complete removal of cohesin from chromosome arms depends on separase. J Cell Sci 2007; 120:4188–96.

57. Misulovin Z, Pherson M, Gause M, Dorsett D. Brca2, Pds5 and Wapl differentially control cohesin chromosome association and function. PLoS Genet 2018; 14:e1007225.

58. Oikawa K, Ohbayashi T, Kiyono T, Nishi H, Isaka K, Umezawa A, Kuroda M, Mukai K. Expression of a novel human gene, human wings apart-like (hWAPL), is associated with cervical carcinogenesis and tumor progression. Cancer Res 2004; 64:3545–9.

59. Losada A. Functional contribution of Pds5 to cohesin-mediated cohesion in human cells and Xenopus egg extracts. J Cell Sci 2005; 118:2133–41.

60. Ohbayashi T, Oikawa K, Yamada K, Nishida-Umehara C, Matsuda Y, Satoh H, Mukai H, Mukai K, Kuroda M. Unscheduled overexpression of human WAPL promotes chromosomal instability. Biochem Biophys Res Commun 2007; 356:699–704.

61. Chude CI, Amaravadi RK. Targeting Autophagy in Cancer: Update on Clinical Trials and Novel Inhibitors. Int J Mol Sci [Internet] 2017 [cited 2018 Sep 16]; 18. Available from: https://www.ncbi.nlm.nih.gov/pmc/articles/PMC5486101/

62. Jiang P, Mizushima N. Autophagy and human diseases. Cell Res 2014; 24:69–79.

63. Simonetti G, Bruno S, Padella A, Tenti E, Martinelli G. Aneuploidy: Cancer strength or vulnerability? Int J Cancer 2018;

64. Vessoni AT, Filippi-Chiela EC, Menck CF, Lenz G. Autophagy and genomic integrity. Cell Death Differ 2013; 20:1444–54.

65. White E, DiPaola RS. The Double-edged Sword of Autophagy Modulation in Cancer. Clin Cancer Res Off J Am Assoc Cancer Res 2009; 15:5308–16.

66. Crasta K, Ganem NJ, Dagher R, Lantermann AB, Ivanova EV, Pan Y, Nezi L, Protopopov A, Chowdhury D, Pellman D. DNA breaks and chromosome pulverization from errors in mitosis. Nature 2012; 482:53–8.

67. Rello-Varona S, Lissa D, Shen S, Niso-Santano M, Senovilla L, Mariño G, Vitale I, Jemaá M, Harper F, Pierron G, et al. Autophagic removal of micronuclei. Cell Cycle Georget Tex 2012; 11:170–6.

68. Bartsch K, Knittler K, Borowski C, Rudnik S, Damme M, Aden K, Spehlmann ME, Frey N, Saftig P, Chalaris A, et al. Absence of RNase H2 triggers generation of immunogenic micronuclei removed by autophagy. Hum Mol Genet 2017; 26:3960–72.

69. Hatch EM, Fischer AH, Deerinck TJ, Hetzer MW. Catastrophic Nuclear Envelope Collapse in Cancer Cell Micronuclei. Cell 2013; 154:47–60.

70. Hintzsche H, Hemmann U, Poth A, Utesch D, Lott J, Stopper H. Fate of micronuclei and micronucleated cells. Mutat Res Mutat Res 2017; 771:85–98.

71. Klionsky DJ, Abdelmohsen K, Abe A, Abedin MJ, Abeliovich H, Acevedo Arozena A, Adachi H, Adams CM, Adams PD, Adeli K, et al. Guidelines for the use and interpretation of assays for monitoring autophagy (3rd edition). Autophagy 2016; 12:1–222.

72. Chung JY-M, Steen JA, Schwarz TL. Phosphorylation-Induced Motor Shedding is Required at Mitosis for Proper Distribution and Passive Inheritance of Mitochondria. Cell Rep 2016; 16:2142–55.

