## Supplementary Figures with Legends for "Lysosomal degradation ensures accurate chromosomal segregation to prevent genomic instability"

**Almacellas et al. Supplemental Information**

### SUPPLEMENTARY FIGURES

**Figure S1.** Related to Figure 1

**Figure S2.** Related to Figure 2

**Figure S3.** Related to Figure 3

**Figure S4.** Related to Figure 4

**Figure S5.** Related to Figure 5

**Figure S6.** Related to Figure 6

### SUPPLEMENTARY VIDEOS

**Video S1.** Related to Figure 1

**Video S2.** Related to Figure 2

**Video S3.** Related to Figure 2

**Video S4.** Related to Figure 2

**Video S5.** Related to Figure 2

**Video S6.** Related to Figure 3

**Video S7.** Related to Figure S3

### SUPPLEMENTARY FIGURES

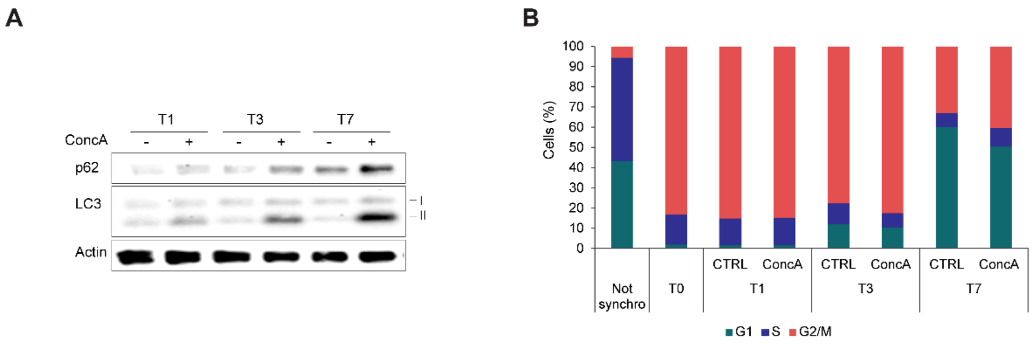


**Figure S1**. (**A**) Time course analysis of U2OS synchronized cells released with or without ConcA (10 nM). LC3 and p62 protein levels were tested at T1, T3 and T7 (1, 3 and 7 hours after RO3306 release, respectively). β-actin protein level was used as loading control. (**B**) Cell cycle analysis of non-synchronized and synchronized U2OS cells treated or not with 10 nM ConcA for the indicated times. Percentage of cells in G1, S and G2/M are represented.

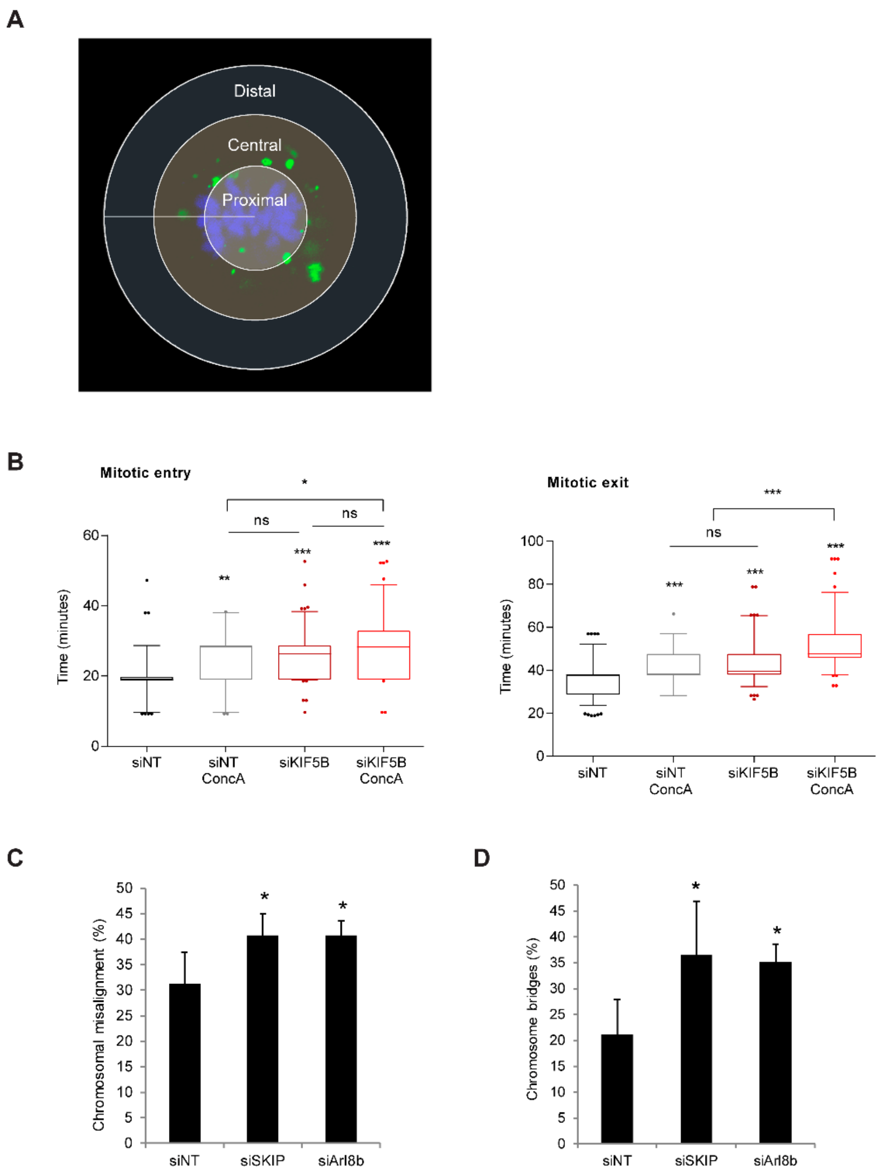


**Figure S2**. (**A**) Representative image of the intracellular distribution of lysosomes in 3 zones: proximal (from center (0) to 100 pixels), central (from 100 to 200 pixels) and distal (from 200 to 300 pixels) using Radial Profile Angle plugin from ImageJ. (**B**) Quantification of mitotic timing of H2B-GFP U2OS cells under the indicated experimental conditions was performed from prophase to metaphase (mitotic entry) or metaphase to cytokinesis (mitotic exit). Error bars represent 5-95 percentiles of 55 ≤ mitosis. (**C-D**) Quantification of mitotic errors (Chromosome misalignment (**C**) and chromosome bridges (**D**) in control U2OS cells or cells depleted for BORC-associated proteins SKIP and Arl8b for 48h using siRNA. Error bars represent S.D of n > 3 experiments Statistical significance is represented as: * $p < 0.05$, ** $p < 0.005$, *** $p < 0.001$ and ns: $p > 0.05$.

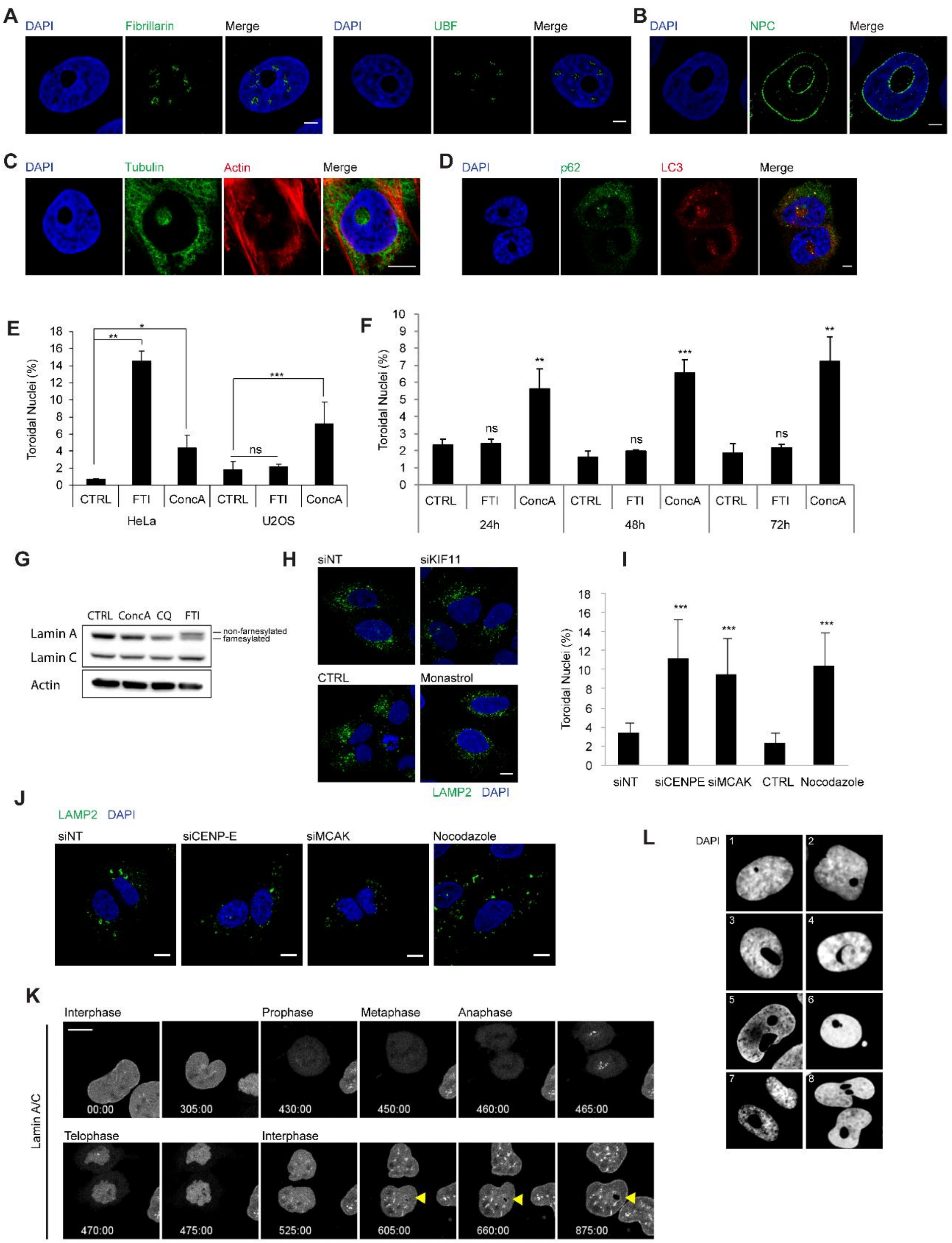


**Figure S3**. (**A-D**) U2OS cells were fixed and processed for immunofluorescence. Single confocal plan image is shown. (**A**) Chromatin-free structure does not correspond to enlarged nucleolus. U2OS cells were fixed and immunofluorescence was performed using fibrillarin or UBF as nucleolar markers and DAPI for DNA detection. Scale bars, 5 µm. (**B**) U2OS cell harboring a toroidal nucleus stained by DAPI for DNA detection (blue) and stained with nuclear pore complex (NPC) antibody to label nuclear membrane (green). Scale bar, 5 µm. (**C**) Detection of phalloidin-positive actin fibers (red) and microtubules (green) demonstrate the presence of cytoskeleton within toroidal nucleus. (**D**) LC3-positive autophagic vesicles and p62 accumulation within toroidal nucleus clarify the presence of cytosolic vesicles through the atypical nucleus. DAPI-stained nuclei in blue. Scale bars, 10 μm. (**E**) Hela and U2OS cells were treated with 10 μM L744832 (FTI) or 10 nM ConcA for 72 h. Cells were fixed and DNA was stained with DAPI for detection of toroidal nuclei. Error bars represent S.D of $n = 3$ experiments (10 fields / experiment). (**F**) U2OS cells were treated with 10 μM L744832 (FTI) or 10 nM ConcA for 24 h, 48 h or 72 h. DAPI-stained DNA was performed to quantify the population of toroidal nuclei compared to total cells. Error bars represent S.D of $n = 3$ experiments (10 fields / experiment). (**G**) U2OS cells were treated with 10 μM L744832 (FTI) or 10 nM ConcA for 72 h and proteins were extracted and analyzed by Western-Blot. Lamin A and Lamin C protein levels were detected. The lower band corresponds to farnesylated Lamin A and the upper ban marks non-farnesylated protein. β-actin was used as loading control. (**H**) Representative single plan confocal images of cells treated with Monastrol for 24 h or depleted for KIF11 for 48 h. Immunofluorescence was performed to detect endogenous lysosomes (LAMP2-positive vesicles in green). Nuclei were stained with DAPI (blue). Scale bar, 10 µm. (**I**) Quantification of the frequency of cells with toroidal nucleus in U2OS cells whether transfected with siRNA control (siNT), siCENP-E, siMCAK for 48 hours or treated or not with microtubule destabilizing agent Nocodazole (1 μM for 24 hours). Error bars represent S.D of $n > 3$ experiments (10 fields / experiment). (**J**) Representative images of single focal plan from experiment described in panel I are shown. Endogenous LAMP2-positive lysosomes are detected by immunofluorescence (green) and nuclei stained with DAPI (blue). Scale bar, 10 µm. (**K**) Live imaging of H2B-GFP mCherry-LaminA/C stable U2OS cell undergoing mitosis for 16 hours every 5 minutes. Nuclear envelope reformation can be observed and precedes toroidal nucleus appearance. Scale bar, 5 µm. (**L**) Single focal plan images of DAPI-stained nuclei (grayscale) in U2OS cells showed the versatility of toroidal nucleus phenotype. Statistical significance is represented as: * $p < 0.05$, ** $p < 0.005$, ***$p < 0.001$, ns: $p > 0.05$.

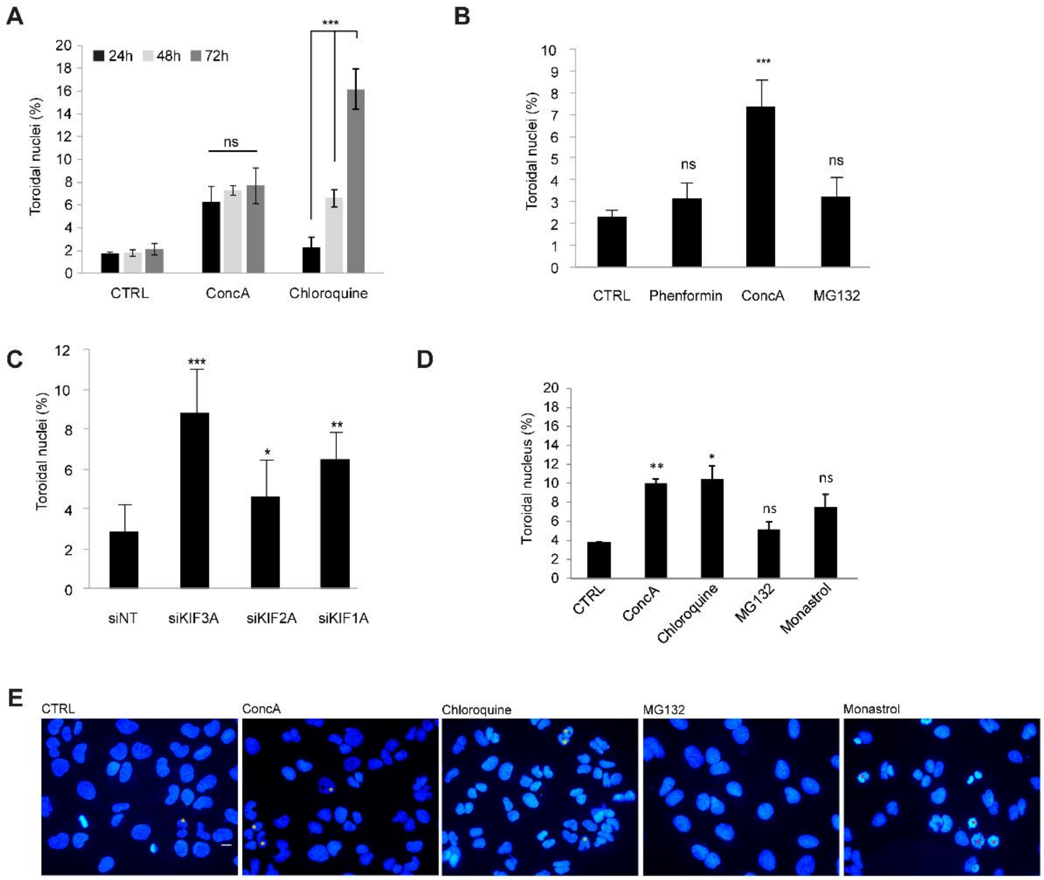


**Figure S4**. (**A**) Toroidal nucleus quantification of U2OS cells treated with ConcA (10 nM) or Chloroquine (10 μM) for the indicated time points and DAPI staining was performed for nucleus detection. Error bars represent S.D. of n > 2 experiments (10 fields / experiment). (**B**) U2OS cells were treated for 24 hours with normal media (CTRL), Phenformin (0.5 mM), ConcA (10 nM) or MG132 (10 μM) and further stained with DAPI for toroidal nucleus frequency quantification. Error bars represent S.D of n > 4 experiments (10 fields / experiment). (**C**) U2OS cells were transfected with siRNA control (siNT), targeting KIF3A (siKIF3A), KIF2A (siKIF2A) or KIF1A (siKIF1A) for 48 hours under normal growth conditions. Nuclei were stained with DAPI for toroidal nucleus quantification. Error bars represent S.D of n ≥ 3 experiments (10 fields / experiment). (**D**) G2/M-synchronized U2OS cells were released in normal media (CTRL) or with 10 nM ConcA, 10 mM Chloroquine, 10 μM MG132 or 100 μM Monastrol. Error bars represent S.D. of n > 3 experiments. (**E**) Representative images of synchronized U2OS processed as in panel **D**. DAPI-stained nuclei are marked with yellow asterisks and cells halted in prometaphase are spotted with red asterisks. Scale bar, 10 μm. **Panels A-D**. Statistical significance is represented as: * $p < 0.05$, ** $p < 0.005$, ***$p < 0.001$, ns: $p > 0.05$.

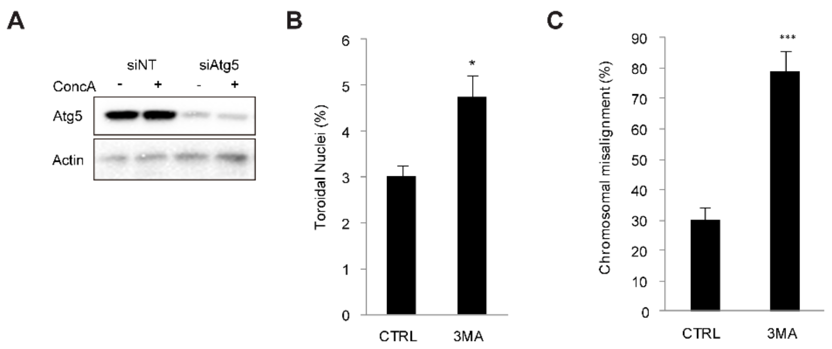


**Figure S5**. (**A**) Validation of Atg5 knockdown (siAtg5) efficiency by Western Blot. β-actin was used as loading control. (**B**) G2/M synchronized cells released with normal media or with 10 mM 3MA for 5 hours. Nuclei were stained with DAPI and toroidal nuclei were quantified. Error bars represent S.D. of n=2 experiments. (**C**) Quantification of chromosomal misalignment in U2OS synchronized cells treated with 3MA (10 mM). Error bars represent S.D. of n=3 experiments (>500 cells). **Panels B** and **C**. Statistical significance is represented as: * $p < 0.05$, ** $p < 0.005$, *** $p < 0.001$, ns: $p > 0.05$.

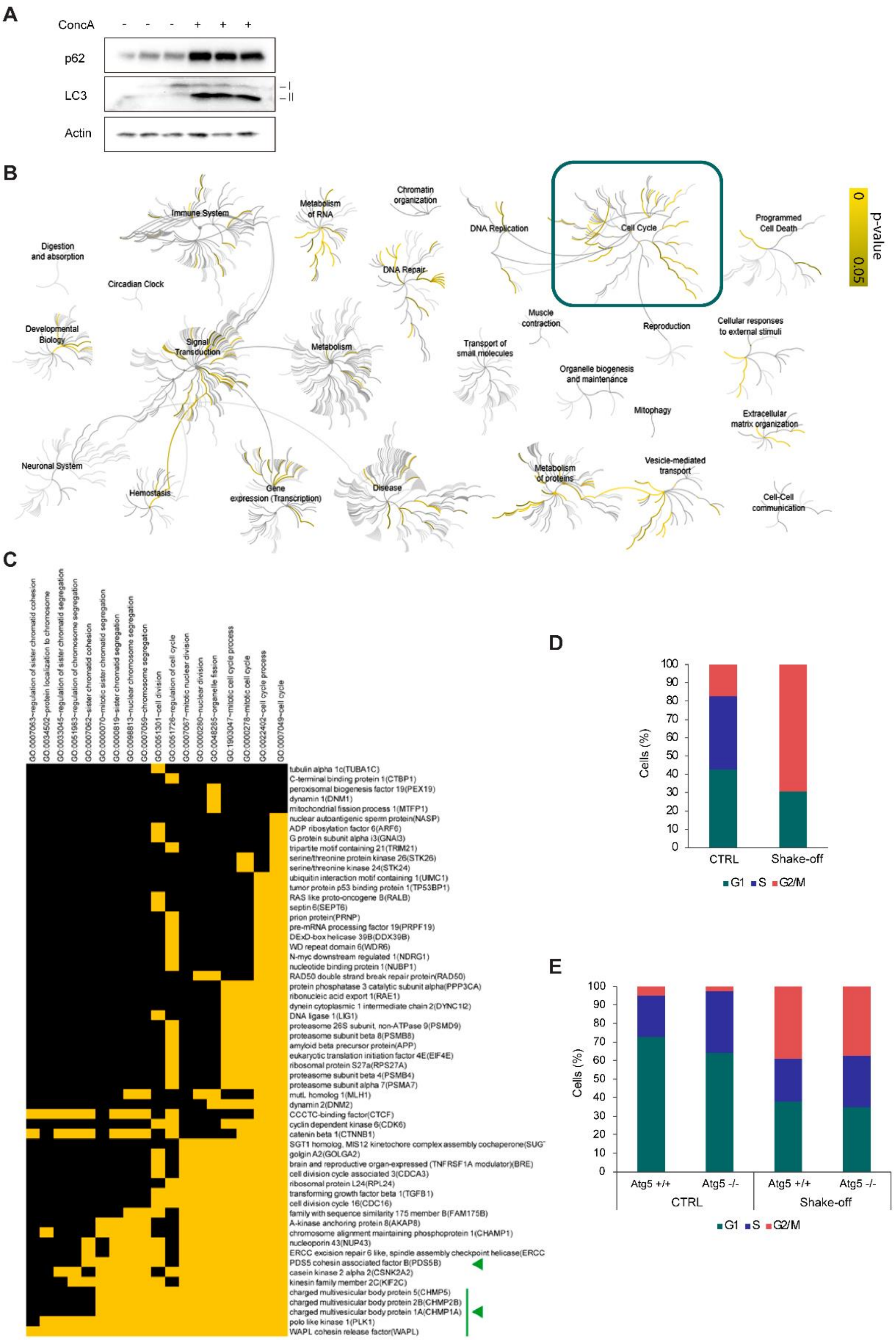
A
ConcA
- - - + + +
p62
LC3
I
II
Actin
B
Immune System
Metabolism of RNA
Chromatin organization
DNA Replication
Cell Cycle
Programmed Cell Death
p-value
0
0.05
Digestion and absorption
Circadian Clock
DNA Repair
Muscle contraction
Reproduction
Cellular responses to external stimuli
Developmental Biology
Signal Transduction
Metabolism
Transport of small molecules
Organelle biogenesis and maintenance
Mitophagy
Extracellular matrix organization
Neuronal System
Hemostasis
Gene expression (Transcription)
Disease
Metabolism of proteins
Vesicle-mediated transport
Cell-Cell communication
C
GO:0007063~regulation of sister chromatid cohesion
GO:0034502~protein localization to chromosome
GO:0033045~regulation of sister chromatid segregation
GO:0051983~regulation of chromosome segregation
GO:0007062~sister chromatid cohesion
GO:0000070~mitotic sister chromatid segregation
GO:0000819~sister chromatid segregation
GO:0098813~nuclear chromosome segregation
GO:0007059~chromosome segregation
GO:0051301~cell division
GO:0051726~regulation of cell cycle
GO:0007067~mitotic nuclear division
GO:0000280~nuclear division
GO:0048285~organelle fission
GO:1903047~mitotic cell cycle process
GO:0000278~mitotic cell cycle
GO:0022402~cell cycle process
GO:0007049~cell cycle
tubulin alpha 1c(TUBA1C)
C-terminal binding protein 1(CTBP1)
peroxisomal biogenesis factor 19(PEX19)
dynamin 1(DNM1)
mitochondrial fission process 1(MTFP1)
nuclear autoantigenic sperm protein(NASP)
ADP ribosylation factor 6(ARF6)
G protein subunit alpha i3(GNAI3)
tripartite motif containing 21(TRIM21)
serine/threonine protein kinase 26(STK26)
serine/threonine kinase 24(STK24)
ubiquitin interaction motif containing 1(UIMC1)
tumor protein p53 binding protein 1(TP53BP1)
RAS like proto-oncogene B(RALB)
septin 6(SEPT6)
prion protein(PRNP)
pre-mRNA processing factor 19(PRPF19)
DExD-box helicase 39B(DDX39B)
WD repeat domain 6(WDR6)
N-myc downstream regulated 1(NDRG1)
nucleotide binding protein 1(NUBP1)
RAD50 double strand break repair protein(RAD50)
protein phosphatase 3 catalytic subunit alpha(PPP3CA)
ribonucleic acid export 1(RAE1)
dynein cytoplasmic 1 intermediate chain 2(DYNC1I2)
DNA ligase 1(LIG1)
proteasome 26S subunit, non-ATPase 9(PSMD9)
proteasome subunit beta 8(PSMB8)
amyloid beta precursor protein(APP)
eukaryotic translation initiation factor 4E(EIF4E)
ribosomal protein S27a(RPS27A)
proteasome subunit beta 4(PSMB4)
proteasome subunit alpha 7(PSMA7)
mutL homolog 1(MLH1)
dynamin 2(DNM2)
CCCTC-binding factor(CTCF)
cyclin dependent kinase 6(CDK6)
catenin beta 1(CTNNB1)
SGT1 homolog, MIS12 kinetochore complex assembly cochaperone(SUGT
golgin A2(GOLGA2)
brain and reproductive organ-expressed (TNFRSF1A modulator)(BRE)
cell division cycle associated 3(CDCA3)
ribosomal protein L24(RPL24)
transforming growth factor beta 1(TGFB1)
cell division cycle 16(CDC16)
family with sequence similarity 175 member B(FAM175B)
A-kinase anchoring protein 8(AKAP8)
chromosome alignment maintaining phosphoprotein 1(CHAMP1)
nucleoporin 43(NUP43)
ERCC excision repair 6 like, spindle assembly checkpoint helicase(ERCC
PDS5 cohesin associated factor B(PDS5B)
casein kinase 2 alpha 2(CSNK2A2)
kinesin family member 2C(KIF2C)
charged multivesicular body protein 5(CHMP5)
charged multivesicular body protein 2B(CHMP2B)
charged multivesicular body protein 1A(CHMP1A)
polo like kinase 1(PLK1)
WAPL cohesin release factor(WAPL)
D
Cells (%)
100
90
80
70
60
50
40
30
20
10
0
CTRL
Shake-off
G1
S
G2/M
E
Cells (%)
100
90
80
70
60
50
40
30
20
10
0
Atg5 +/+
Atg5 -/-
Atg5 +/+
Atg5 -/-
CTRL
Shake-off
G1
S
G2/M

**Figure S6**. (**A**) Western Blot analysis of synchronized U2OS cell fractions (2) processed for mass spectrometry analysis. Validation of ConcA efficiency was assessed by p62 and LC3-I and LC3-II protein levels. β actin was used as loading control. (**B**) Representation of pathways affected by lysosome impairment during cell division. Results from mass spectrometry were subjected to functional annotation analysis using online Reactome tool (https://reactome.org/). Yellow lines represent significantly affected pathways ($p$-value $< 0.05$). Green square highlights altered cell cycle processes. (**C**) Functional annotation clustering of proteins related to cell cycle regulation and mitotic progression. Mass spectrometry results were filtered and protein hits that presented a fold change of more than 1.2 (Conc A vs ctrl) and those only present in Conc A cells were analyzed using David GO gene ontology web tool. Functional annotation clustering was performed. The heat map represents all the candidate proteins involved in cell cycle regulation and mitotic progression. Yellow squares indicate matching with the GO term described; Black squares for missing relation. (**D**) Cell cycle analysis of U2OS cellular fractions obtained by shake-off or under normal growing condition (CTRL). Propidium iodide staining was used to follow the DNA. The percentage of cells in each cell cycle phase (G1, S, G2/M) is represented. (**E**) Cell cycle analysis of cellular fractions from MEF Atg5 wt and KO cells growing in normal conditions (CTRL) or obtained by shake-off. Propidium iodide stained DNA. The percentage of cells in each cell cycle phase (G1, S, G2/M) is represented.

### SUPPLEMENTARY VIDEOS

**Video 1**. Lysosomes are dynamic during cell division. U2OS cells stably expressing H2B-GFP (blue) and LAMP1-RFP (arbitrary in green) were subjected to live imaging with confocal microscope every 5 minutes for 24 hours. Scale bar, 10 μm. Supporting video of **Figure 1A**.

**Videos 2-4**. Mitotic delay induced by lysosome impairment. Supporting videos of **Figure 2F**.

**Video 5**. Toroidal nucleus formation. Supporting video of **Figure 3B**.

**Video 6**. Imaris animation. Supporting video of **Figure 3F**.

**Video 7**. Nuclear envelop reformation. Supporting video of **Supplementary Figure 3E**.
