## Supplementary material for "Lysosomal degradation ensures accurate chromosomal segregation to prevent genomic instability": Material and Methods Tables

| Chemicals | Working Concentration | Catalogue Number |
|---|---|---|
| Concanamycin A | 10 nM | Santa Cruz SC-20211A |
| Chloroquine | 10 μM | SIGMA C6628 |
| L744832 (FTI) | 10 μM | MERCK 422720 |
| Monastrol | 100 μM | SIGMA M8515 |
| Nocodazole | 1 μM | SIGMA M1404 |
| MG132 | 10 μM | SIGMA M7449 |
| RO3306 | 9 μM | SIGMA SML0569 |
| Thymidine | 25 μM | SIGMA T1895 |
| Phenformin | 0.5 mM | SIGMA P7045 |

**Table 1**. List of the chemicals used in this study.

| Gene | Sequence [dT][dT] (5' - 3') | [siRNA] | Brand / Gift from |
|---|---|---|---|
| ARL8B | GAUAGAAGCUUCCCGAAAU | Dharmacon | J-020294-09-0005 |
| ATP6V0c | GGCACAGCCAAGAGCGGUA<br>GCUCUGUGUAUGCGGAUGA<br>CCCGACUAUUCGUGGGCAU<br>CCAGCUAUCUAUAACCUUA | Dharmacon | L-017620-01-0005 |
| Atg5 | CATCTGAGCTACCCGGATA | 50 nM | gift from Dr. Muñoz (IDIBELL) |
| CENP-E | GGAAUUAAAGGCUAAAAGA | 20 nM | SIGMA |
| KIF1A | GGAAACAGAGAAGAUCAUA | 20 nM | SIGMA |
| KIF2A | GAAAUGGUUUACAGGUUUA<br>CUACACAACUUGAAGCUAU<br>GAAAACGACCACUCAAUAA<br>GACCCUCCUUCAAGAGAUA | Dharmacon | L-004959-00 |
| KIF2C/MCAK | GCAAGCAACAGGUGCAAGU | 20 nM | SIGMA |
| KIF3A | GGUGUUCGAGCUAUUCCUG | 20 nM | SIGMA |
| KIF5B | CGGCGACAAGUACAUCGCCAAGUUU | 20 nM | SIGMA |
| KIF11 | GCUACUCUGAUGAAUGCAU | 20 nM | SIGMA |
| p62/SQSTM1 | GCAUUGAAGUUGAUAUCGAU | 50 nM | gift from Dr. Muñoz (IDIBELL) |
| SKIP | CUUCUGAACUGGACCGAUU | 20 nM | SIGMA |

**Table 2**. List of siRNA used in this study.

| Primary antibodies | | | |
|---|---|---|---|
| Antigen | Dilution | Source | Catalog Number |
| ATG5 | WB 1/1000 | Abgent | AP1812b |
| ß-Actin | WB 1/10000 | Sigma | A2228 |
| ß-Actin | WB 1/2000 | Cell Signaling | 4967 |
| Fibrillarin | IF 1/75 | Santa Cruz | sc-25397 |
| LC3 (pAb) | WB 1/1000, IF 1/100 | MBL | pm036 |
| LAMP-2 (CD107b) | IF 1/300 | BD Biosciences | 555803 |
| LAMIN A/C | WB 1/1000 | CST | 2032S |
| LAMIN B1 | IF 1/100 ON | Abcam | ab16048 |
| NPC | IF 1/100 ON | Abcam | Ab-24609 |
| Nucleolin | IF 1/100 | Santa Cruz | sc-13057 |
| p62 / SQSTM1 | WB 1/1000, IF 1/200 | MBL | M162-3 |
| PDS5B | WB 1/1000 | Bethyl Lab. | A300-537A-M |
| Pericentrin | IF 1/100 | Abcam | ab28144 |
| Phalloidin - Rhodamine | IF 1/500 | Cytoskeleton | Phdh1 |
| α-tubulin | IF 1/250 | Sigma-Aldrich | T6074 |
| UBF | IF 1/100 ON | Santa Cruz | sc-13125 |
| WAPL | WB 1/1000 | Bethyl Lab. | A300-268A-M |
| Secondary antibodies | | | |
| Antigen | Dilution | Source | Catalog Number |
| Anti-Mouse Ig HRP | WB 1/5000 | Dako | P0260 |
| Anti-Rabbit Ig HRP | WB 1/5000 | Dako | P0399 |
| Anti-Mouse (goat), IgG (H+L), Alexa Fluor 488 | IF 1/400 | Invitrogen | A11001 |
| Anti-Rabbit (goat), IgG, (H+L), Rhodamine Red-X | IF 1/400 | Invitrogen | R6394 |
| Anti-Rabbit (goat), IgG (H+L), Alexa Fluor 546 | IF 1/400 | Thermofisher | A11010 |
| Anti-Rabbit (donkey), IgG, (H+L), Alexa Fluor 555 | IF 1/400 | Thermofisher | A31572 |

**Table 3**. List of primary and secondary antibodies used in this study.
